# Closing the conformational cycle of the beta-barrel assembly machinery

**DOI:** 10.64898/2026.10.20.760484

**Authors:** Takaaki Kato, Hideki Shigematsu, Koh Takeuchi, Masato Suzuki, Ryo Nitta, Tsuyoshi Imasaki

## Abstract

The β-barrel assembly machinery (BAM) catalyzes the folding and insertion of β-barrel outer membrane proteins (OMPs) in Gram-negative bacteria without an external energy source. BAM comprises the central subunit BamA and associated lipoproteins, with BamA forming a conformationally dynamic lateral gate. Although substrate-engaged structures have revealed key steps in OMP assembly, how substrate-free BamA closes its lateral gate and which conformations it adopts in a lipid bilayer remain unclear. Here, we determined cryo-EM structures of the *Escherichia coli (E.coli)* BAM complex in three detergents and in Saposin A nanodiscs composed of *E. coli* polar lipids. In addition to the canonical outward-open (OO) state, we identified an occluded (OC) state in which both the lateral gate and the periplasmic opening of the BamA barrel are closed. OC predominated in lipid-reconstituted BAM that retained OMP-folding activity, indicating that this state is substantially populated in a functional BAM ensemble within a lipid bilayer. We also identified a semi-occluded (sOC) state in which β1–β2 adopts an OC-like closed position while rearrangement of β16 remains incomplete. Comparison of OO, sOC, and OC reveals asynchronous rearrangements of the β1–β2 and β16 regions, with β1–β2 movement can occur without complete repositioning of β16. Re-examination of previously reported lipid-embedded BAM structures further highlights the importance of local map quality at the gate region for conformational assignment. Together, our findings expand the BamA conformational landscape beyond the conventional OO–inward-open (IO) framework to include OC and sOC as distinct structural states associated with lateral-gate closure following OMP insertion.

## Introduction

The outer membrane of Gram-negative bacteria is an asymmetric lipid bilayer with lipopolysaccharide (LPS) in its outer leaflet and forms a formidable permeability barrier that limits the entry of many compounds, including antibiotics. Its biogenesis depends on numerous β-barrel outer membrane proteins (OMPs), whose folding and insertion are catalyzed by the essential β-barrel assembly machinery (BAM)^1,2^. In *Escherichia coli*, BAM consists of the central subunit BamA and four associated lipoproteins, BamB–E, and functions without using an external energy source such as ATP. BamA is essential for bacterial viability and target of antibacterial molecules including darobactin^3,4^. BamA contains a 16-stranded transmembrane β-barrel and five periplasmic POTRA domains. A lateral gate formed between β1 and β16 undergoes large conformational changes that have been broadly described as outward-open (OO) and inward-open (IO) states ^5–7^.

Structures of substrate-engaged BAM have revealed key steps in OMP assembly, including β-augmentation between BamA and nascent substrates, hybrid-barrel formation, and progressive folding of the substrate β-barrel^8–15^. In contrast, the structural events that follow substrate insertion remain less well understood^16^. It is unclear how the BamA lateral gate closes after substrate release, whether closure occurs through a concerted motion or additional intermediate states, and which conformation predominates in substrate-free BAM. These questions are especially relevant because opening of the lateral gate is thought to locally thin and destabilize the surrounding membrane, thereby facilitating OMP insertion^5,10,17^. A more tightly sealed conformation may therefore be favored after substrate release, but such a state has not been clearly defined.

Most structural analyses of BAM have relied on purified complexes examined in detergents or lipid-based membrane mimetics, where its conformational landscape may depend on the surrounding membrane environment and on the dynamics of peripheral subunits. Substrate-free BAM structures determined under these conditions have been assigned to different lateral-gate states^8,17–19^, but conformational heterogeneity and limited local resolution around β1 and β16 have complicated direct comparison. Moreover, detergents and lipid mimetics can alter the relative stability of membrane-protein conformations, yet systematic comparisons of the same BAM cross distinct membrane environments remain limited. BamB and the other accessory subunits may also contribute additional structural heterogeneity, raising the question of whether peripheral-subunit dynamics are directly coupled to BamA gating or instead represent a separate conformational variable.

Here, we biochemically characterized and structurally compared *E. coli* BAM in three detergents, DDM, GDN, and LMNG, and in Saposin A nanodiscs. Cryo-EM analysis revealed the canonical OO state in DDM, an occluded (OC) state in GDN and lipid nanodiscs, and a semi-occluded (sOC) state in LMNG. In OC, both the lateral gate and the periplasmic opening of BamA are closed, whereas sOC exhibits an OC-like displacement of β1–β2 with incomplete rearrangement of β16. The lipid-reconstituted BAM retained OMP-folding activity, and independent analyses showed that BamB ordering varied markedly with membrane environment but was only weakly coupled to BamA gate state. Together, these results define a broader conformational landscape of substrate-free BAM and suggest that lateral-gate closure involves asynchronous, separable rearrangements of the two strands of the β1–β16 seam.

## Results

### BAM maintains a common subunit architecture across membrane-mimetic conditions

We purified the *E. coli* BAM complex (BamABCDE) and reconstituted it in one of the following detergents: n-dodecyl-β-D-maltoside (DDM), glyco-diosgenin (GDN), or lauryl maltose neopentyl glycol (LMNG), or reconstituted it into Saposin A nanodiscs containing *E. coli* polar lipid extract. Under all four conditions, BAM was purified from a monodisperse peak by size-exclusion chromatography (Figure 1a).

**Figure 1.**
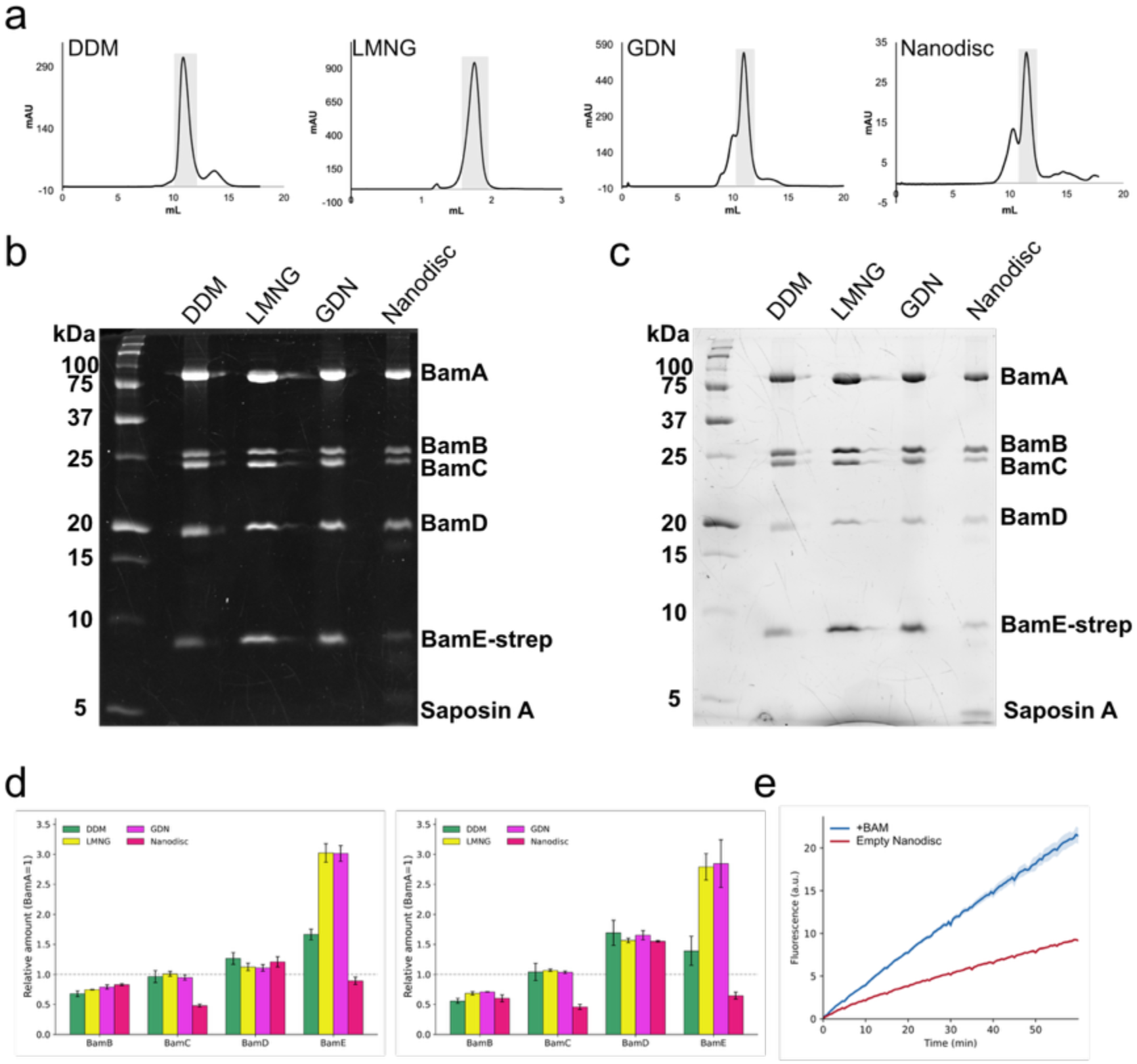
Purification and subunit composition of BAM under the four conditions. (a) Size-exclusion chromatography profiles for BAM in DDM, LMNG, GDN, and Nanodisc. Gray boxes highlight the main elution peaks. (b) Stain-free (TCE-based, Trp-dependent) image of the SDS-PAGE gel, with subunit bands labeled. (c) CBB-stained image of the same gel. (d) Relative amounts of BamB to BamE (BamA = 1), corrected by Trp number for stain-free (left) and by molecular weight for CBB (right). Bars are colored by condition and error bars show the standard deviation (N = 3). (e) Folding activity measured as fluorescence over 60 min for BAM-containing nanodiscs (BAM, blue) and empty nanodiscs (EMP, red). Lines show the mean and shaded bands the standard deviation (n = 3).

To assess whether the membrane-mimetic conditions altered subunit composition, we analyzed the same samples using two independent gel-based detection methods, stain-free^20^ detection (Figure 1b) and Coomassie Brilliant Blue (CBB) staining (Figure 1c). Band intensities were corrected for tryptophan content for stain-free detection or molecular mass for CBB staining and expressed as apparent relative abundance normalized to BamA (N = 3; Figure 1d).

Across all four conditions, BAM retained all five subunits (BamA–E) with broadly similar apparent subunit compositions (Figure 1d). The relative abundances of BamB, BamC, and BamD were comparable across all conditions, with the exception of a lower apparent abundance of BamC in saposin A nanodiscs. Apparent BamE abundance exceeded that of BamA under detergent conditions, with stain-free BamE/BamA ratios of 1.66 in DDM, 3.02 in LMNG, and 3.01 in GDN, whereas the ratio was 0.89 in Saposin A nanodiscs. Because BAM is expected to contain one copy of each subunit, the elevated BamE signal likely reflects condition-dependent effects of purification and quantification rather than a true change in complex stoichiometry. The use of BamE-Strep for affinity capture may contribute to this apparent enrichment, particularly during the more extensive detergent-exchange washes used for LMNG and GDN.

We next tested whether BAM reconstituted into Saposin A nanodiscs retained OMP-folding activity using an OmpT, a β-barrel outer membrane protease and substrate of BAM. BAM-containing nanodiscs exhibited substantially greater OmpT-dependent fluorescence than empty nanodiscs, reflecting successful OmpT folding and demonstrating that the reconstituted BAM population remained functional (Figure 1e; Figure S1).

Together, these biochemical analyses showed that BAM remained intact under all four membrane-mimetic conditions and that the lipid-reconstituted sample retained OMP-folding activity, providing a basis for subsequent structural comparisons.

### Cryo-EM reveals membrane-environment-dependent conformations of BamA

We next determined single-particle cryo-EM structures of BAM under all four membrane-mimetic conditions (Figure 2; Figures S2–S4; Table 1). All five subunits (BamA–E) were present in every final reconstruction, indicating that the major particle populations retained the intact BAM architecture. The core subunits BamA, BamC, BamD, and BamE were consistently well resolved, whereas BamB was present in all maps but showed pronounced condition-dependent differences in structural ordering.

**Figure 2.**
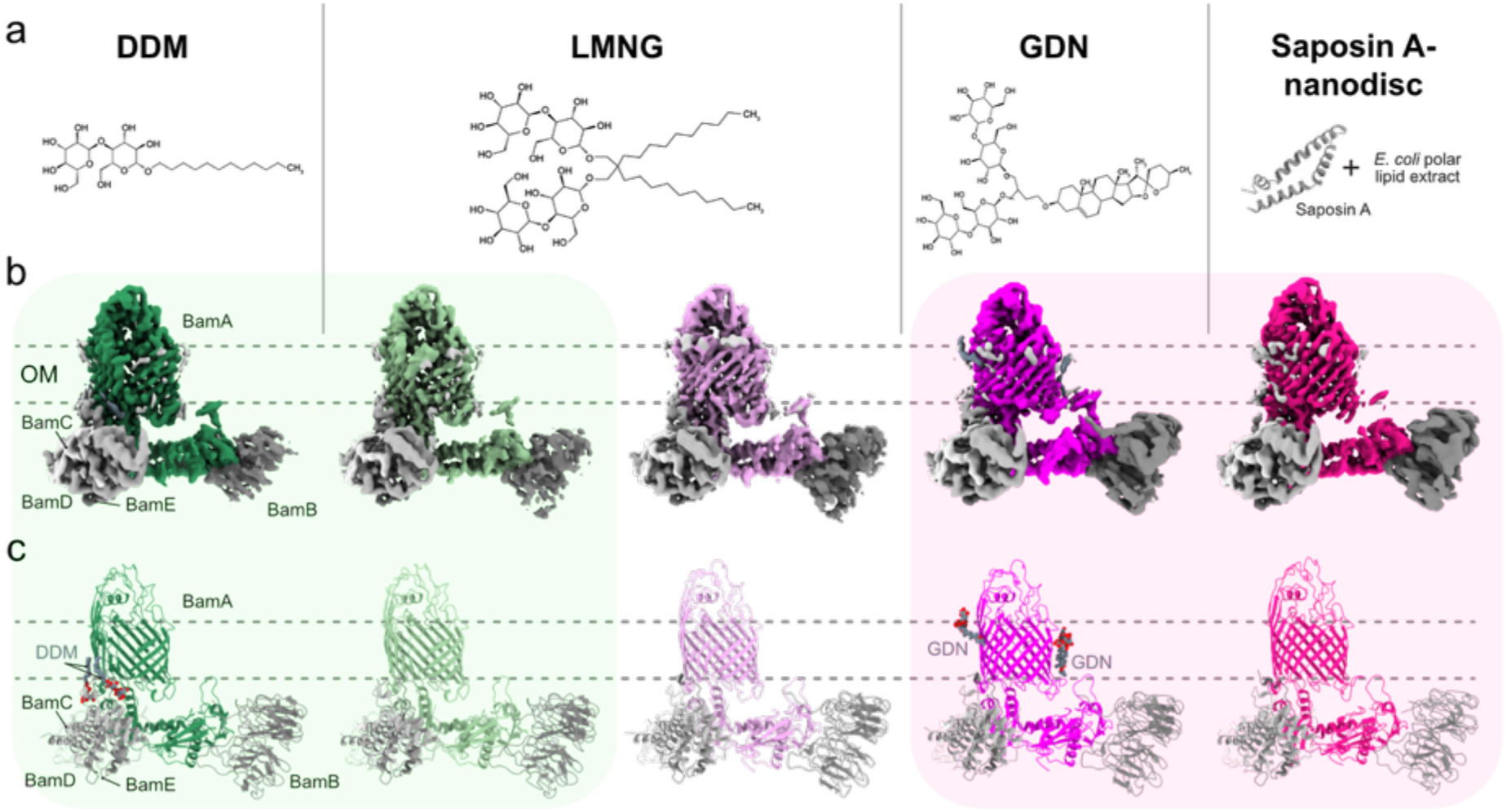
Cryo-EM analysis of the *E. coli* BAM under four membrane-mimetic conditions. (a) Chemical structures of the three detergents DDM, LMNG, and GDN, and a schematic of the Saposin A nanodisc reconstituted with *E. coli* polar lipid extract. (b) Cryo-EM density maps of BAM in DDM, LMNG, GDN, and Saposin A nanodisc. For LMNG, two conformations are shown. BamA is colored and the peripheral subunits BamB to BamE are shown in shades of gray. The approximate position of the outer membrane (OM) is indicated by dashed lines. (c) Atomic models corresponding to the maps in (b), colored as in (b). Bound DDM and GDN molecules are shown as spheres.

**Table 1.**
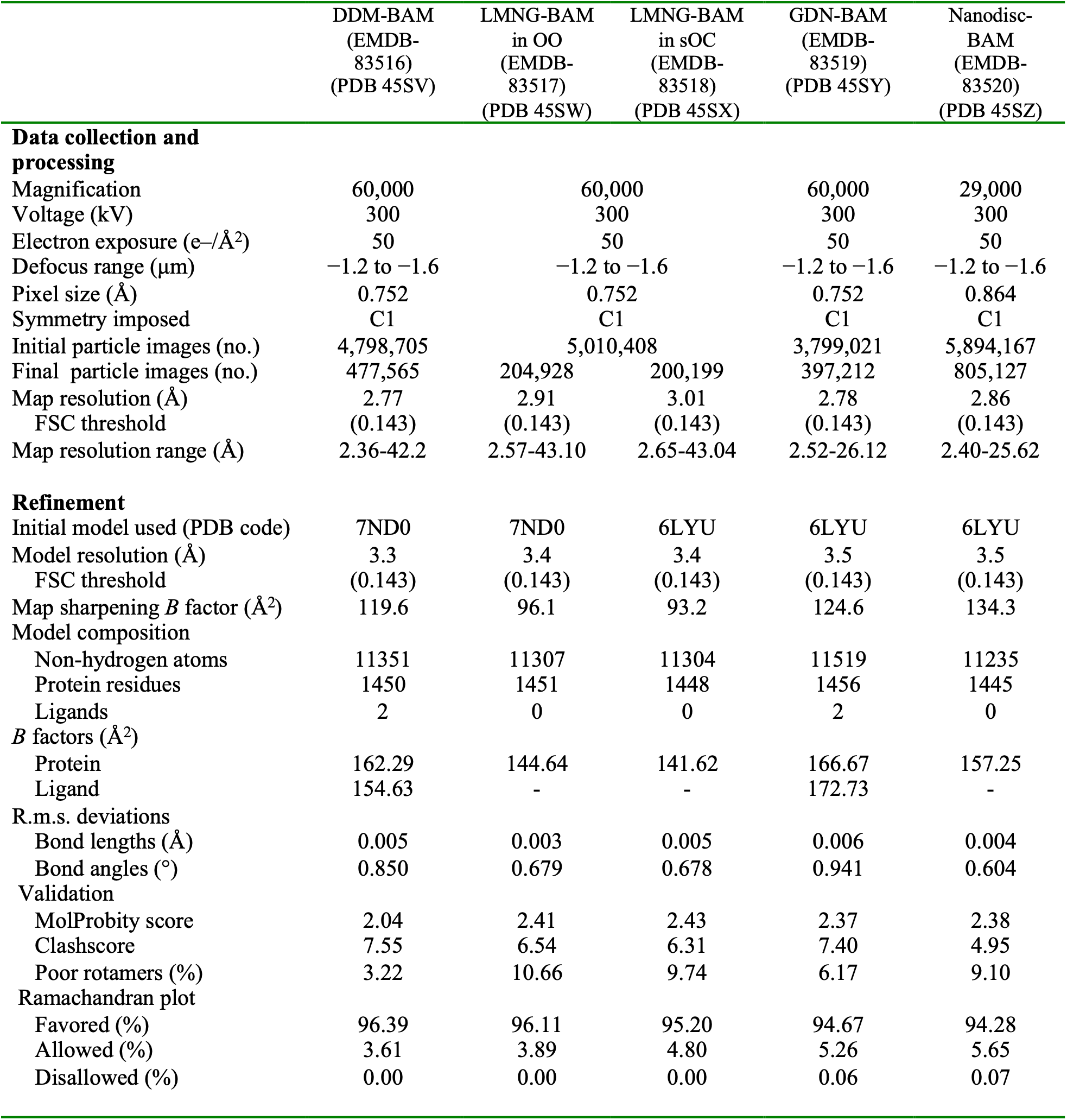
Cryo-EM data collection, refinement and validation statistics.

Rigid-body comparison of the structures showed that most of the BAM was conserved, whereas the largest conformational differences were concentrated around the lateral gate of the BamA β-barrel (Figure S5). DDM predominantly stabilized an outward-open (OO) conformation, GDN and Saposin A nanodiscs predominantly stabilized a closed conformation, and LMNG supported two distinct gate states. We therefore used the DDM structure as a reference to define the membrane-environment-dependent conformational changes described below.

### DDM stabilizes the outward-open state of BamA

We determined the BAM structure in DDM micelles at an overall resolution of 2.77 Å (Figure 3; green). The BamA lateral gate was widely open, with the β1–β2 region containing extracellular loop 1 (eL1) displaced from β16. The structure closely matched previously reported outward-open (OO) BAM structures (RMSD = 0.84 Å to PDB 9CNW)^19^, including the positions of β1–β2, β16, and eL1. We therefore assigned the DDM structure as the canonical OO state and used it as a reference for subsequent comparisons^6,21,22^.

**Figure 3.**
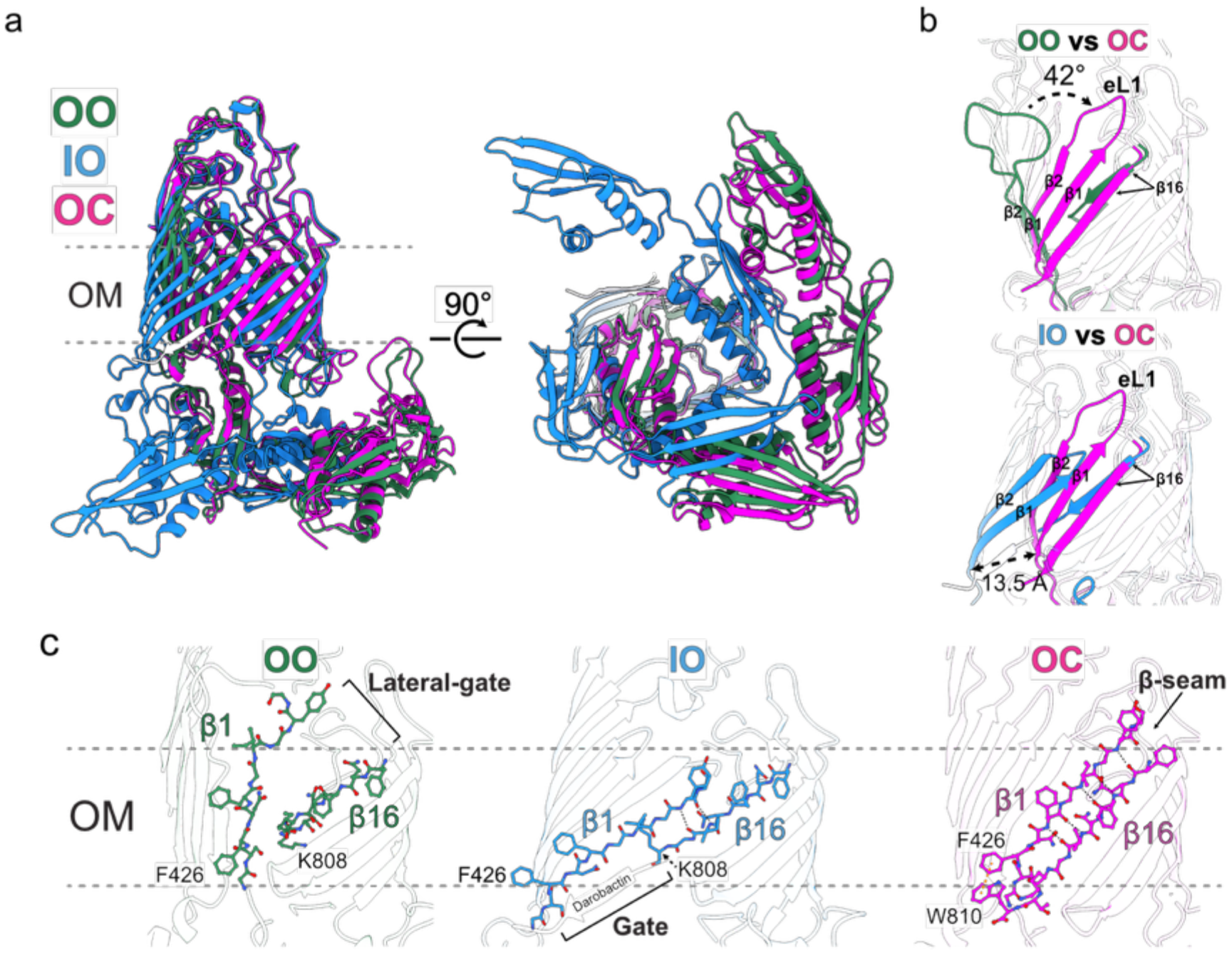
GDN-reconstituted BAM adopts a previously undefined occluded state with both gates closed. (a) Superposition of the outward-open (OO, green), inward-open (IO, blue), and occluded (OC, magenta) states of BamA, in two views related by a 90° rotation. (b) Close-up of the lateral gate. In the OO versus OC comparison (top), the eL1-containing β1-β2 region rotates by about 42°. In the IO versus OC comparison (bottom), the F426 (β1 strand) is displaced by about 13.5 Å. (c) Lateral-gate region of the OO, IO, and OC states, with β1, β16, and F426 indicated. The gate, the darobactin bound to the IO state, and the β-seam of the OC state are labeled.

Two DDM molecules could be modeled into well-defined densities along hydrophobic grooves on the transmembrane surface of the BamA β-barrel (Figure S6a). These molecules were distributed along the barrel surface rather than being selectively concentrated at the lateral gate.

### GDN reveals an occluded state with closed lateral and periplasmic openings

We determined the BAM structure in GDN micelles at an overall resolution of 2.78 Å (Figure 3; magenta). Well-resolved GDN molecules were also observed along hydrophobic surfaces of the BamA transmembrane β-barrel (Figure S6b). Relative to DDM-OO, the β1–β2 region, including eL1, rotated by approximately 42° toward β16, closing the lateral gate (Figure 3a, b). GDN-BAM was also distinct from the darobactin-bound inward-open (IO) state^4^. Superposition with IO showed a displacement of approximately 13.5 Å at the Cα atom of F426 at the base of β1.

The two structures also differed on the periplasmic side. Whereas the barrel lumen is accessible from the periplasm in IO, POTRA domain 5 (P5) covers the periplasmic opening in GDN-BAM (Figure S7). Thus, both the lateral gate and the periplasmic opening are closed. We therefore define this conformation, which is distinct from both OO and IO, as the occluded (OC) state.

At the closed lateral gate, β1 and β16 pair to form a β-seam. F426 on β1 also packs closely against W810 near the C terminus of β16, with an aromatic ring-centroid distance of approximately 4.6 Å (Figure 3c; Figure S3). The well-resolved side-chain densities support this interaction, identifying F426–W810 packing as a characteristic feature of OC.

### LMNG captures a semi-occluded gate conformation

BAM in LMNG displayed pronounced heterogeneity at the lateral gate. Focused 3D classification of the β1–β3 region separated the particles into two nearly equally populated classes. One closely matched DDM-OO (RMSD = 0.65 Å; Figure 4a; Table S2) and was assigned as LMNG-OO.

**Figure 4.**
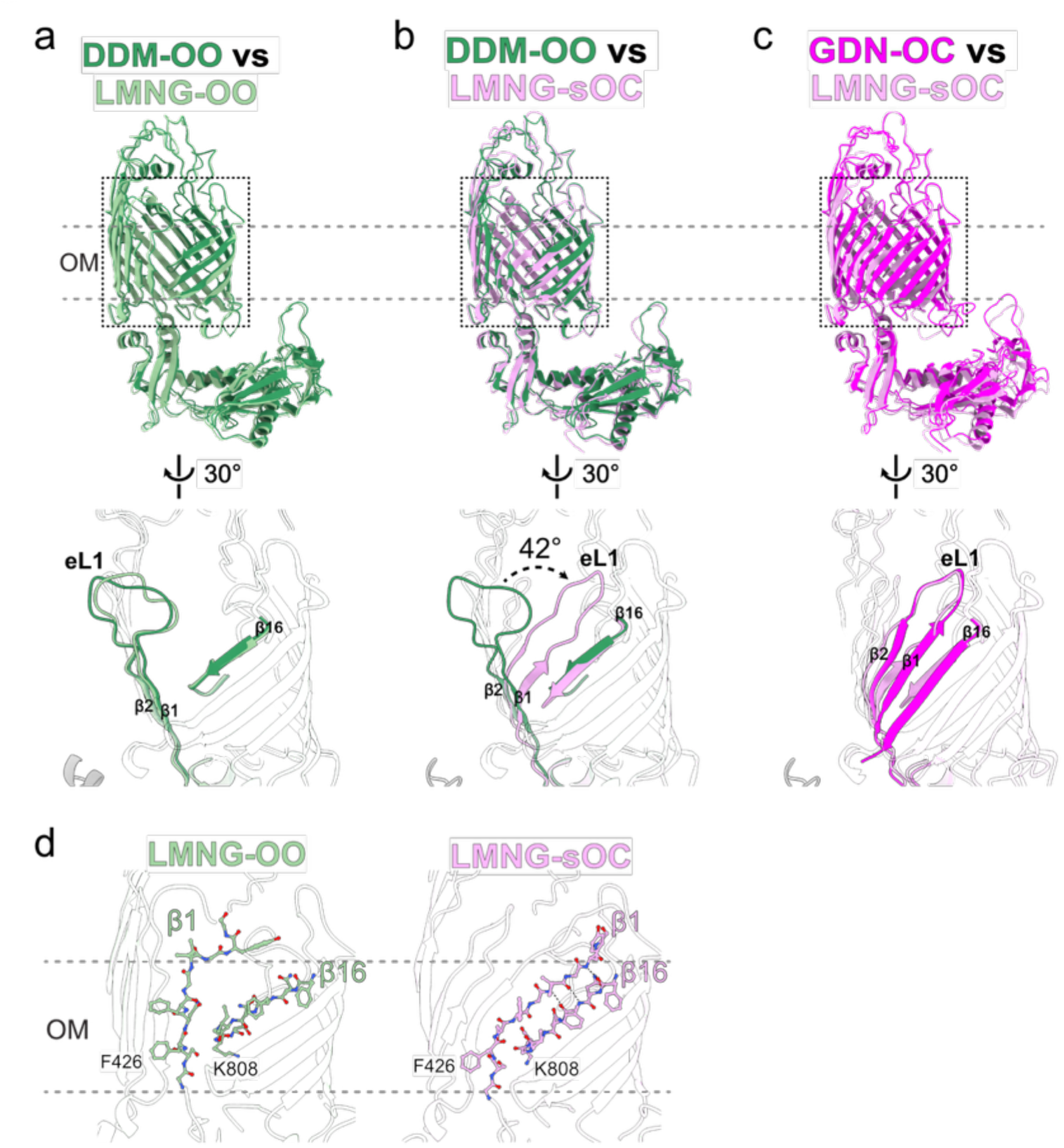
LMNG reveals a semi-occluded intermediate between the OO and OC states. (a) Superposition of DDM-OO and LMNG-OO, with a close-up of the lateral gate below. (b) Superposition of DDM-OO and LMNG-sOC. The eL1-containing β1-β2 region of LMNG-sOC rotates by about 42° toward β16. (c) Superposition of GDN-OC and LMNG-sOC. (d) Lateral-gate region of LMNG-OO and LMNG-sOC, with β1, β16, F426, and T7 indicated.

In the second class, β1–β2 rotated by approximately 42° toward β16 and adopted an OC-like position (Figure 4b, c). β16, however, remained displaced from its position in OC, leaving the β-seam incomplete and preventing formation of the F426–W810 interaction. Thus, movement of the β1–β2 side of the gate was largely complete, whereas rearrangement of the β16 side remained incomplete. These changes were confined to the lumenal side of the gate, without affecting the periplasmic pore. We define this conformation as the semi-occluded (sOC) state.

To assess the robustness of the classification, we randomly divided the particle set into two halves and independently repeated focused 3D classification. Both subsets reproduced OO and sOC at similar proportions (Figure S8), supporting sOC as a reproducible BamA conformation that coexists with OO in LMNG.

### The occluded state predominates in functional BAM embedded in a lipid bilayer

We next examined the cryo-EM structure of the functional BAM sample reconstituted into Saposin A nanodiscs containing *E. coli* polar lipid extract. The structure was determined at an overall resolution of 2.86 Å (Figures S2 and S3; Table 1).

The predominant nanodisc structure closely matched GDN-OC (RMSD = 1.9 Å; Figure 5a; Table S2). The lateral gate was closed, β1 and β16 formed the β-seam, the F426–W810 interaction was present, and P5 covered the periplasmic opening of the barrel (Figure 5b).

**Figure 5.**
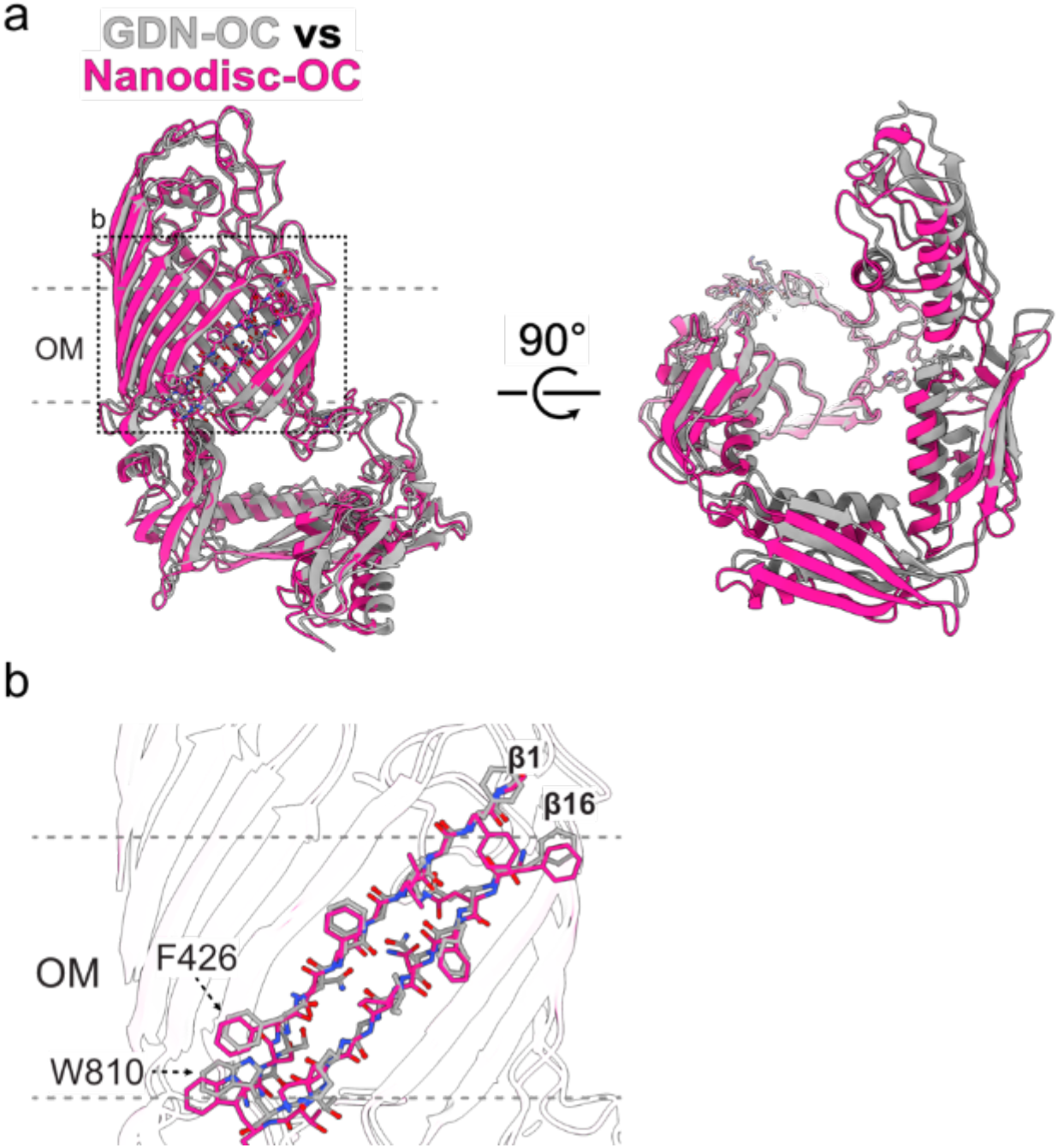
The occluded conformation is preserved in a lipid bilayer with a well-resolved lateral gate. (a) Superposition of GDN-OC (gray) and nanodisc-OC (magenta), in two views related by a 90° rotation. (b) Close-up of the lateral gate, showing the β1–β16 pairing and the F426–W810 interaction. β1, β16, F426, and W810 are indicated.

Thus, OC is not restricted to BAM solubilized in GDN but also predominates in a lipid-bilayer environment. Because biochemical analysis established that the nanodisc-reconstituted BAM population retains OMP-folding activity, OC represents a major conformation accessible to functional lipid-embedded BAM. However, the OmpT folding assay measures the activity of the complete conformational ensemble and therefore does not establish that OC itself directly catalyzes OmpT folding.

### BAMA gate conformation and BamB ordering are weakly coupled

In addition to altering BamA gate conformation, the membrane-mimetic environment markedly affected BamB ordering. cryoDRGN^23,24^ analysis estimated that the fraction of particles with weakly ordered BamB decreased from 51% in DDM and 38% in LMNG to 15% in GDN and 5% in Saposin A nanodiscs (Figure S9a, b). These differences were substantially greater than the differences in apparent BamB abundance measured biochemically (less than 20% difference), suggesting that they predominantly reflect changes in BamB ordering or flexibility rather than differences in subunit occupancy.

We independently assessed BamB ordering using LocOccupancy^25^, which evaluates local map occupancy without particle classification. The fraction of BamB residues with local occupancy below 0.8 was 22% in DDM, 23% in LMNG, 16% in GDN, and 12% in Saposin A nanodiscs (Figure S9b). By contrast, local occupancy of the BamA barrel remained uniformly high under all conditions (median ≥ 0.98), arguing against differences in overall map quality as the source of the BamB variation. Although cryoDRGN and LocOccupancy measure different properties and their absolute values are not directly comparable, both analyses independently showed greater BamB disorder in DDM and LMNG and greater ordering in GDN and nanodiscs.

We next asked whether BamB ordering was coupled to the BamA gate conformation. LMNG provided a direct test because OO and sOC coexist under the same biochemical condition. Particles assigned to OO and sOC extensively overlapped in the cryoDRGN latent space (ROC-AUC = 0.53; Figure S9c, d), and the fractions with weakly ordered BamB were similar in the two classes (36.3% and 40.0%, respectively; Cramér’s V = 0.038). Thus, although both BamA and BamB respond to the membrane environment, BamA lateral-gate conformation and BamB ordering are only weakly coupled in LMNG and represent largely separable dimensions of BAM structural heterogeneity.

### Comparison with previously reported BAM structures in lipid environments

Several *E. coli* BAM structures have previously been determined in lipid environments using Saposin A nanodiscs^8^, membrane scaffold protein (MSP)-based nanodiscs^18^, or styrene-maleic acid (SMA)-solubilized native lipids^19^ and have been described as laterally closed or outward-open conformations (Figure S10a).

Comparison with our Saposin A-OC structure showed broadly similar overall BamA architectures but substantial differences in local map quality at the lateral gate. In several previously reported maps, density around β1–β2 and β16, the region most critical for assigning the gate conformation, was relatively weak (Figure S10b). Consistently, Q-score^26^ and DAQ analyses^27^ indicated lower local map-model agreement at the gate, with mean gate Q-scores ranging from 0.28 to 0.41 and 41–65% of gate residues showing negative DAQ values (Figure S11; Table S3). In our reconstructions, by contrast, the lateral gate was resolved at a quality comparable to the remainder of the BamA barrel. The gate Q-scores were 0.70 for SaposinA-OC and 0.61 for DDM-OO, compared with 0.61 and 0.64, respectively, for the remainder of the barrel (gate-to-rest ratios of 1.15 and 0.95). Consistently, the mean gate DAQ scores were positive, at 0.64 and 0.56, respectively, with no gate residues exhibiting negative DAQ values, including those in β16 (Figure S11; Table S3). Thus, both the absolute local quality metrics and their comparison with the remainder of the barrel support reliable interpretation of the lateral-gate region in our reconstructions.

In the Saposin A-OC reconstruction, β1 and β16 were individually resolved, allowing direct assessment of the β-seam and F426–W810 interaction. The β16 C-terminal region, including W810, was also fully resolved and modeled, whereas previously reported bilayer structures (PDB 6LYU, 7RI5, and 9CNY) were modeled only to residue 806.

Together, these observations highlight the importance of local map quality when assigning BamA lateral-gate conformations. The present reconstruction provides a direct structural definition of the OC lateral gate supported by well-resolved density for both sides of the β1–β16 seam.

### OO, sOC, and OC delineate asymmetric rearrangements associated with lateral-gate closure

Comparison of OO, sOC, and OC identified a series of asymmetric rearrangements associated with closure of the BamA lateral gate (Figure 6). Relative to OO, the β1–β2 region in OC rotated by approximately 42° and β3–β4 by approximately 20°, accompanied by rearrangements of β14–β15, periplasmic turn 7 (T7), and β16. Displacement of T7 opened additional space adjacent to F426 on β1, allowing W810 near the C terminus of β16 to move alongside F426 and establish the F426–W810 grip that stabilizes the closed gate. Rearrangements were also observed in the T5–T6 region and the P3 loop on the periplasmic side. In OC, these movements brought W627, V208, V209, and F724 into close proximity, producing tighter packing around the closed gate.

**Figure 6.**
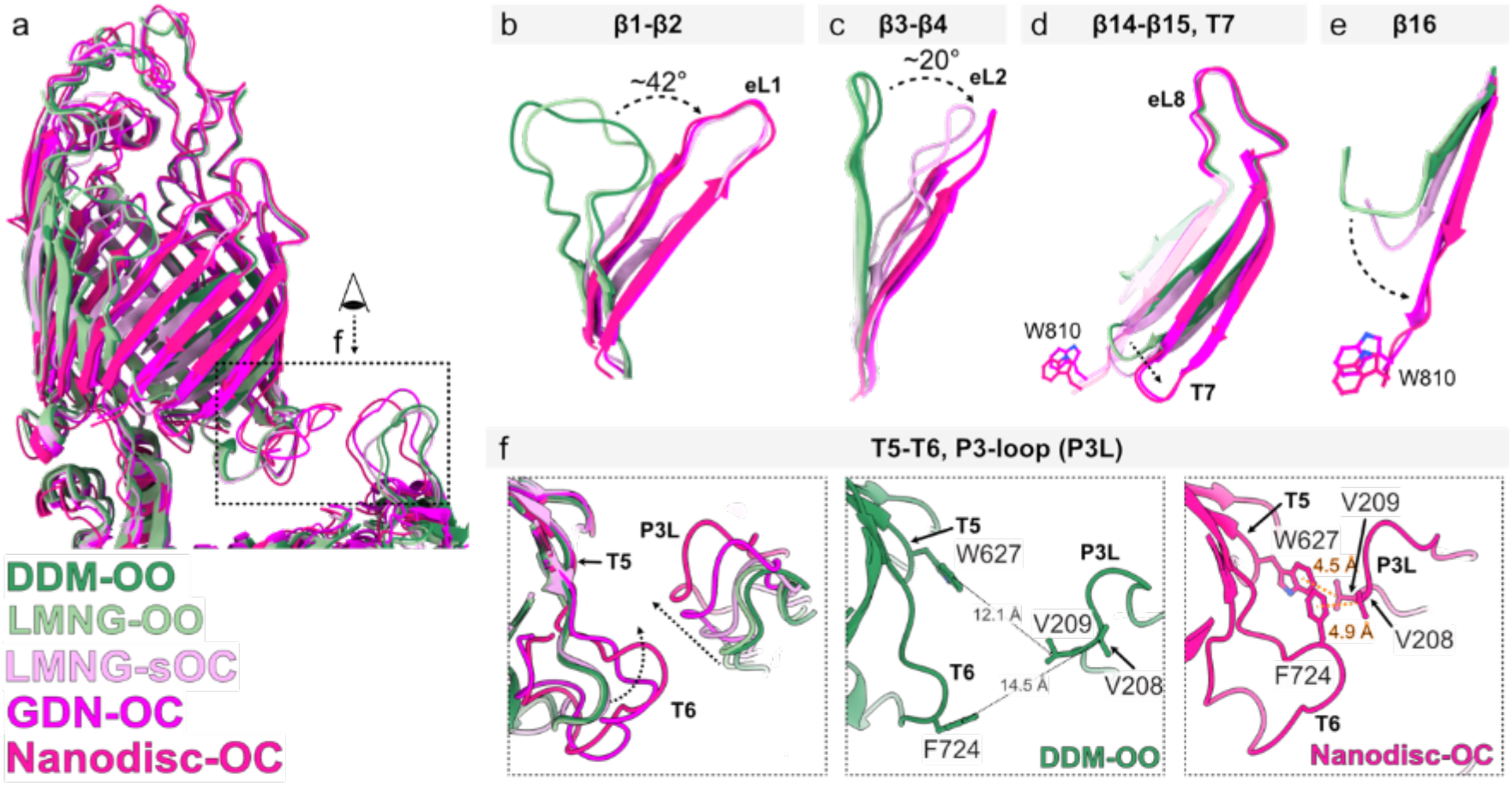
The lateral gate adopts a range of conformations from open to fully sealed. (a) Superposition of DDM-OO, LMNG-OO, LMNG-sOC, GDN-OC, and nanodisc-OC. The boxed region is enlarged in (f). (b-e) State-dependent rearrangements of individual gate elements. β1-β2 rotates by about 42° (b), β3-β4 by about 20° (c), while β14-β15 and T7 (d) and β16 (e) also change position. W810 is indicated in (d) and (e). (f) Periplasmic T5-T6 and the P3 loop (P3L) rearrange between states. In nanodisc-OC, W627, V208, V209, and F724 come into close contact (4.5 and 4.9 Å), whereas the corresponding distances in DDM-OO are larger (12.1 and 14.5 Å).

Distance measurements further resolved the individual components of gate closure (Figure S12; Table S4). The Y432–F802 distance across the β1–β16 gate decreased from 13–15 Å in OO to approximately 6 Å in both sOC and OC. In contrast, the F426–K808 distance, which reports rearrangement on the β16 side, decreased from approximately 11 Å in OO to 8.2 Å in sOC and approximately 5 Å in OC. Correspondingly, the number of backbone hydrogen bonds between β1 and β16 increased from none in OO to three in sOC and nine in OC.

These measurements show that β1–β2 has already reached an almost OC-like position in sOC, whereas β16 remains incompletely rearranged. The structural differences among OO, sOC, and OC therefore indicate that the two sides of the lateral gate can rearrange asynchronously rather than as a single rigid-body movement.

OC was also distinct from IO on the periplasmic side. The periplasmic opening measured approximately 20 Å in OO and OC but expanded to approximately 31 Å in IO (Figure S12; Figure S4) . Consistently, pore-radius analysis showed a minimum radius of approximately 1 Å at the periplasmic opening in both OO and OC, compared with approximately 6.5 Å in IO. Thus, OC is not simply a laterally closed form of IO but a distinct BamA conformation in which both the lateral gate and the periplasmic opening are closed.

## Discussion

Our study expands the conformational landscape of BamA beyond the conventional OO–IO framework. Systematic comparison of the same BAM across membrane-mimetic environments revealed two additional conformations: an OC state in which both the lateral gate and the periplasmic opening are closed, and a sOC state in which β1–β2 has adopted an OC-like position while rearrangement of β16 remains incomplete. Importantly, OC predominated not only in GDN but also in lipid-reconstituted BAM that retained OMP-folding activity, indicating that OC is not restricted to a detergent-stabilized complex but is accessible to functional BAM in a lipid bilayer.

Comparison of OO, sOC, and OC reveals that lateral-gate closure involves structurally separable rearrangements of its two sides (Figure 7). In sOC, β1–β2 has rotated by approximately 42° toward β16, reducing the gate width to nearly that of OC, whereas β16 remains incompletely repositioned. In OC, repositioning of β16 completes the β1–β16 seam and brings F426 and W810 into close contact. The progressive shortening of distances across the gate and formation of backbone hydrogen bonds further support an asymmetric reorganization of the gate. Thus, lateral-gate closure need not occur as a concerted rigid-body motion: the β1 side can reach an essentially closed position while the β16 side remains incompletely repositioned. sOC therefore represents an asymmetric gate configuration rather than simply an average between the open and closed structures. The separability of these rearrangements suggests that partially closed configurations such as sOC can be populated without complete formation of the β1–β16 seam. Although the structural continuity among OO, sOC, and OC suggests a plausible closure pathway, these structures are static snapshots and do not establish their temporal order or interconversion pathway.

**Figure 7.**
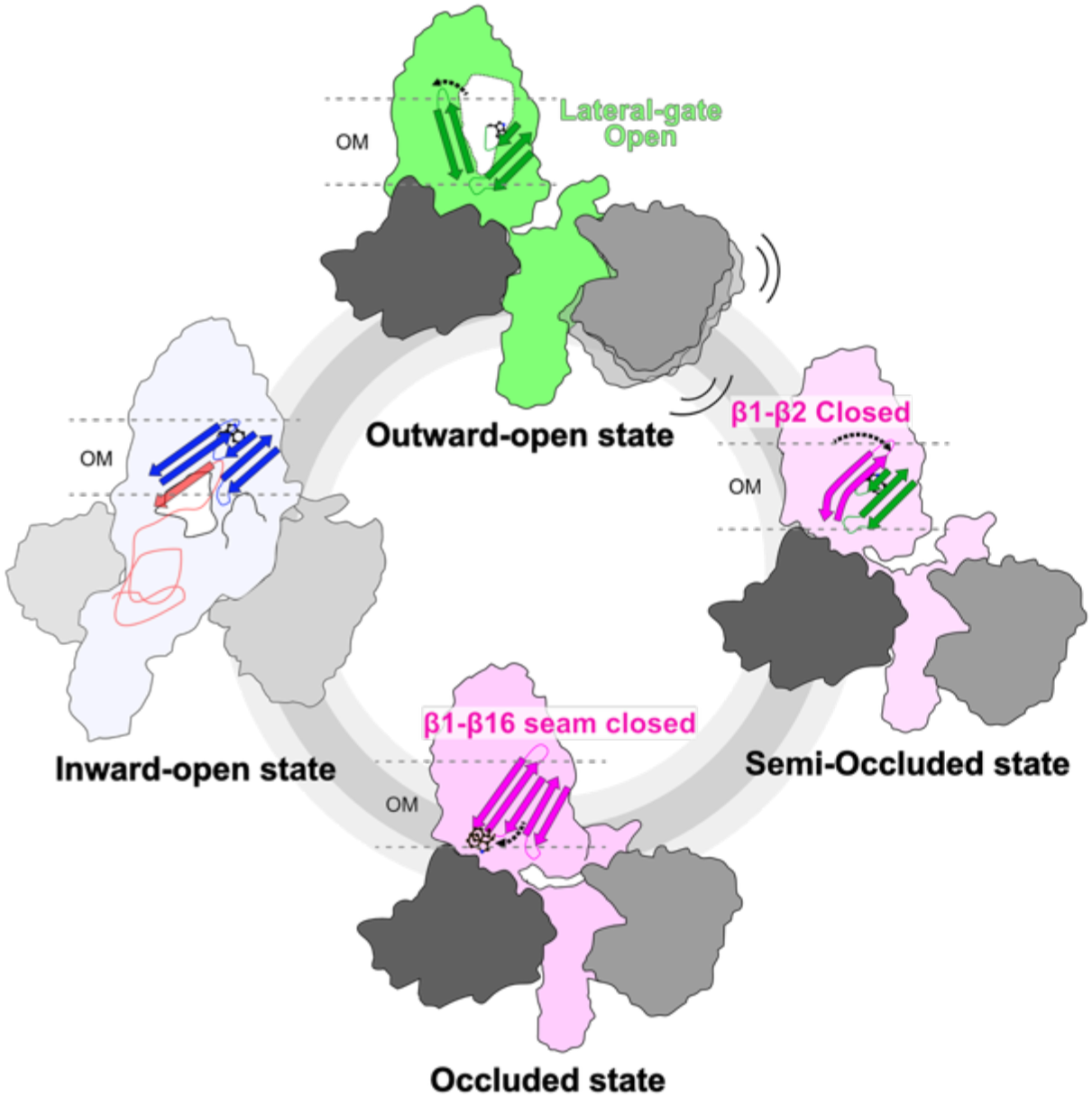
Expanded conformational landscape of BamA and structural model for lateral-gate closure. Schematic model integrating the previously characterized inward-open (IO) and outward-open (OO) states with the semi-occluded (sOC) and occluded (OC) states identified in this study. In sOC, β1–β2 adopts an OC-like closed position, whereas β16 remains incompletely repositioned. In OC, repositioning of β16 completes the β1–β16 seam and brings F426 and W810 into close contact. The arrangement of the four states illustrates the expanded conformational landscape of BamA and does not establish their temporal order or a sequential interconversion pathway.

The architecture of OC suggests a potential role as a substrate-free resting conformation. In OC, the β1–β16 seam seals the lateral gate, while P5 restricts access to the barrel lumen from the periplasm, distinguishing OC from both OO and IO. Notably, β1, which provides the edge for β-augmentation with substrate OMPs during assembly, instead pairs with β16 in OC, and this self-paired configuration is further stabilized by the F426–W810 interaction. These features suggest a simple switching mechanism in which the β1 edge alternates between self-pairing with β16 in a closed state and β-augmentation with substrate during OMP assembly^8–11,14^.

Because this closed configuration would need to remain reversible during the assembly cycle, substrate delivery may promote reopening of the lateral gate. Unfolded OMPs are transferred to BAM by periplasmic chaperones such as SurA, and substrate engagement could therefore favor disruption of the β1–β16 seam and shift BamA toward a substrate-receptive conformation^12,13,15^. In this framework, sOC and OC may represent structural states associated with the return of BamA toward a substrate-free configuration after OMP insertion. Whether OC is the predominant resting state in the native outer membrane, whether sOC lies on the physiological transition pathway, and how substrate binding drives reopening of the gate remain to be established.

The distinct state distributions observed in DDM, LMNG, GDN, and nanodiscs further indicate that the membrane environment reshapes the BamA conformational landscape. DDM favored OO, LMNG supported both OO and sOC, whereas GDN and lipid nanodiscs favored OC. These environments need not create otherwise inaccessible conformations. Rather, they may alter the relative energetic stability of states intrinsically sampled by BamA^28,30^ as illustrated by the closely related OC structures obtained in both GDN and a lipid bilayer. The positions of resolved detergent molecules provide a possible structural basis for some of these differences. In DDM, a resolved detergent molecule occupied a hydrophobic groove at the BamA–BamD interface that overlaps the closed-state position of β16 and W810, potentially favoring the outward-open conformation. In GDN, the resolved detergents occupied peripheral sites compatible with gate closure, whereas no ordered detergent was resolved near the gate in LMNG. Together, these observations suggest that the observed state distributions cannot be readily attributed to a single conserved detergent-binding site and may instead reflect a combination of local interactions and broader properties of the membrane-mimetic environment. Our comparison with previously reported lipid-embedded BAM structures further highlights the importance of local map quality when assigning lateral-gate conformations, particularly for β1, β16, and the C-terminal region of β16. More broadly, systematic comparison across membrane mimetics, combined with local assessment of structural heterogeneity, provides a useful strategy for resolving conformational states of dynamic membrane proteins.

The influence of membrane environment extended beyond BamA gating. BamB was substantially less ordered in DDM and LMNG than in GDN and nanodiscs, a trend independently supported by cryoDRGN and local-occupancy analyses. Because these differences were considerably greater than the variation in apparent BamB abundance measured biochemically, they are more consistent with conformational disorder than with loss of BamB. Moreover, within LMNG, BamB ordering was only weakly associated with whether BamA adopted OO or sOC. Thus, the membrane environment appears to modulate at least two partially separable dimensions of BAM heterogeneity. Our gel-based measurements nevertheless report apparent relative abundance rather than absolute stoichiometry, and additional orthogonal measurements will be required to distinguish changes in subunit occupancy from conformational disorder.

From a bacteriological perspective, a tightly closed substrate-free conformation could help minimize unnecessary membrane perturbation between rounds of OMP assembly, whereas reopening of the lateral gate would permit productive substrate engagement. Perturbation of this conformational balance could compromise OMP biogenesis and outer-membrane integrity, potentially affecting bacterial susceptibility to antibiotics and other environmental stresses. Thus, OC and sOC may represent functionally relevant states associated with maintenance of the bacterial envelope. This expanded conformational landscape may also be relevant to antibacterial inhibition. Darobactin stabilizes an inward-open BamA conformation^4^, whereas other BamA inhibitors exhibit state-dependent binding^19^. The identification of OC and sOC therefore broadens the structural context for considering conformation-selective inhibition of BamA.

Several limitations define the next steps. Saposin A nanodiscs containing *E. coli* polar lipid extract provide a lipid bilayer but do not reproduce the asymmetric architecture and distinct physicochemical environment of the native outer membrane, including its LPS-rich outer leaflet^29, 31^. Moreover, the structures identified here were stabilized under different conditions and do not directly establish their interconversion kinetics, and the OmpT assay demonstrates activity of the ensemble rather than catalytic activity of OC itself. Determining the populations of OO, sOC, OC, and other states in native outer membranes will therefore be important. In situ structural approaches, together with state-selective mutations and kinetic or single-molecule measurements, should test whether the asymmetric closure inferred here occurs during the physiological assembly cycle. Collectively, our findings show that the BamA lateral gate can undergo asymmetric, structurally separable rearrangements and identify OC and sOC as states that expand the conformational landscape beyond the canonical OO–IO framework.

## Supplementary Figures

**Supplementary Figure 1.**
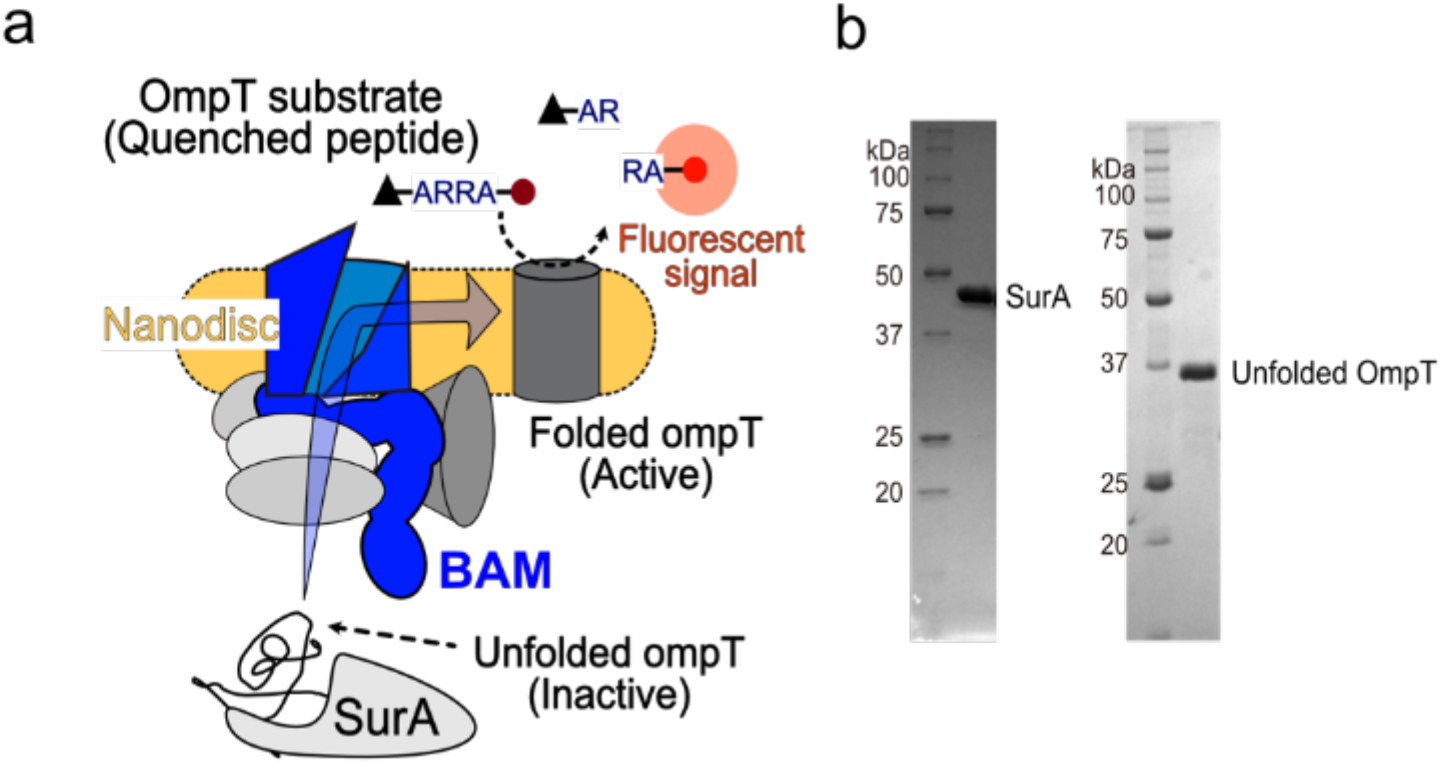
Schematic of the OmpT folding assay and characterization of purified SurA and unfolded OmpT. (a) Schematic of the in vitro folding assay. Unfolded OmpT delivered by the chaperone SurA is folded by BAM reconstituted in a nanodisc. Folded OmpT cleaves a quenched peptide and generates a fluorescent signal. (b) SDS-PAGE of the purified SurA and of the unfolded OmpT substrate used in the assay.

**Supplementary Figure 2.**
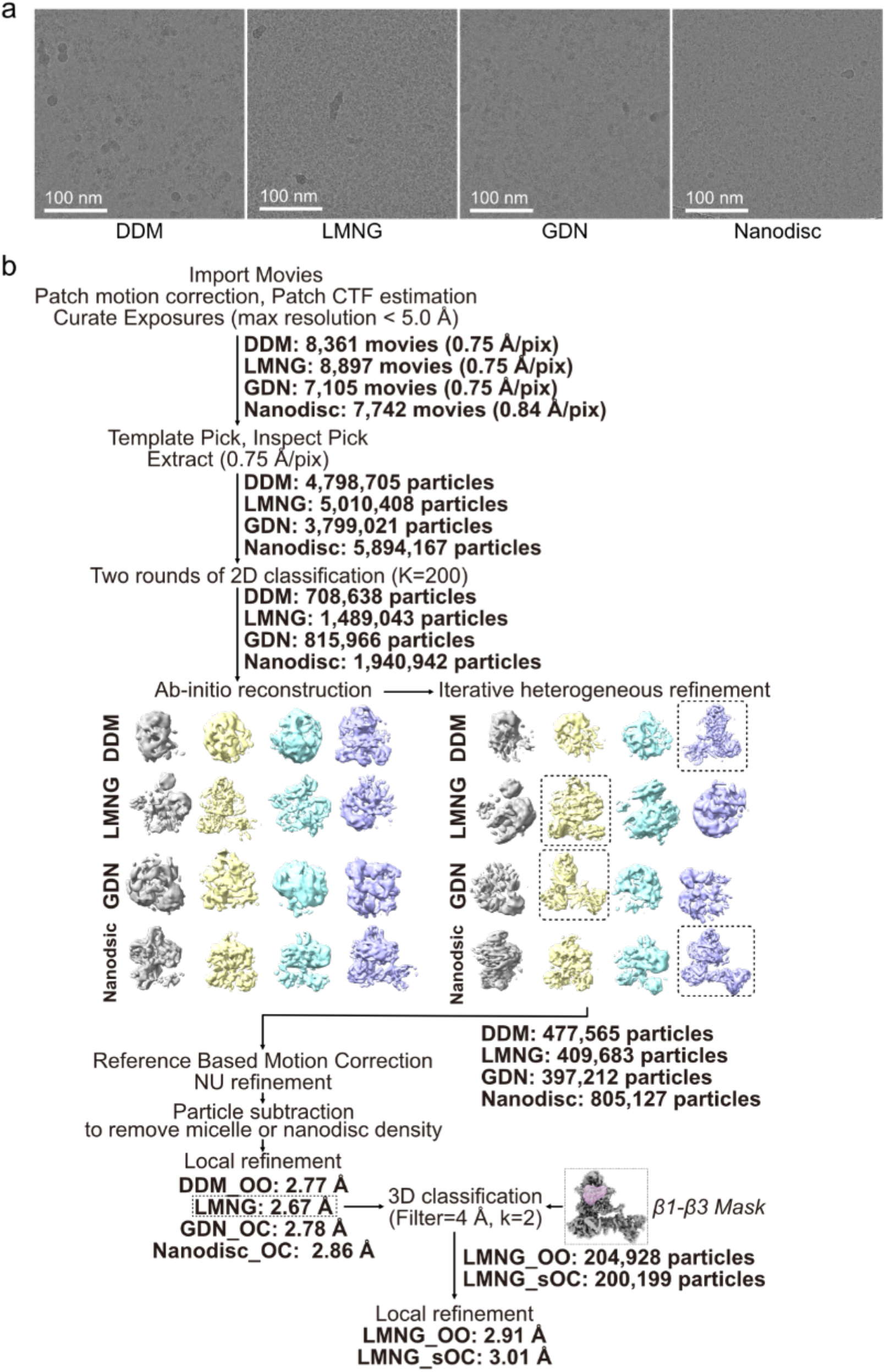
Cryo-EM data acquisition and image processing. (a) Representative motion-corrected micrographs for each condition. Scale bar, 100 nm. (b) Image-processing workflow, including movie and particle numbers, classification steps, and final resolutions. For LMNG, focused 3D classification with a lateral-gate mask separated the OO and sOC classes.

**Supplementary Figure 3.**
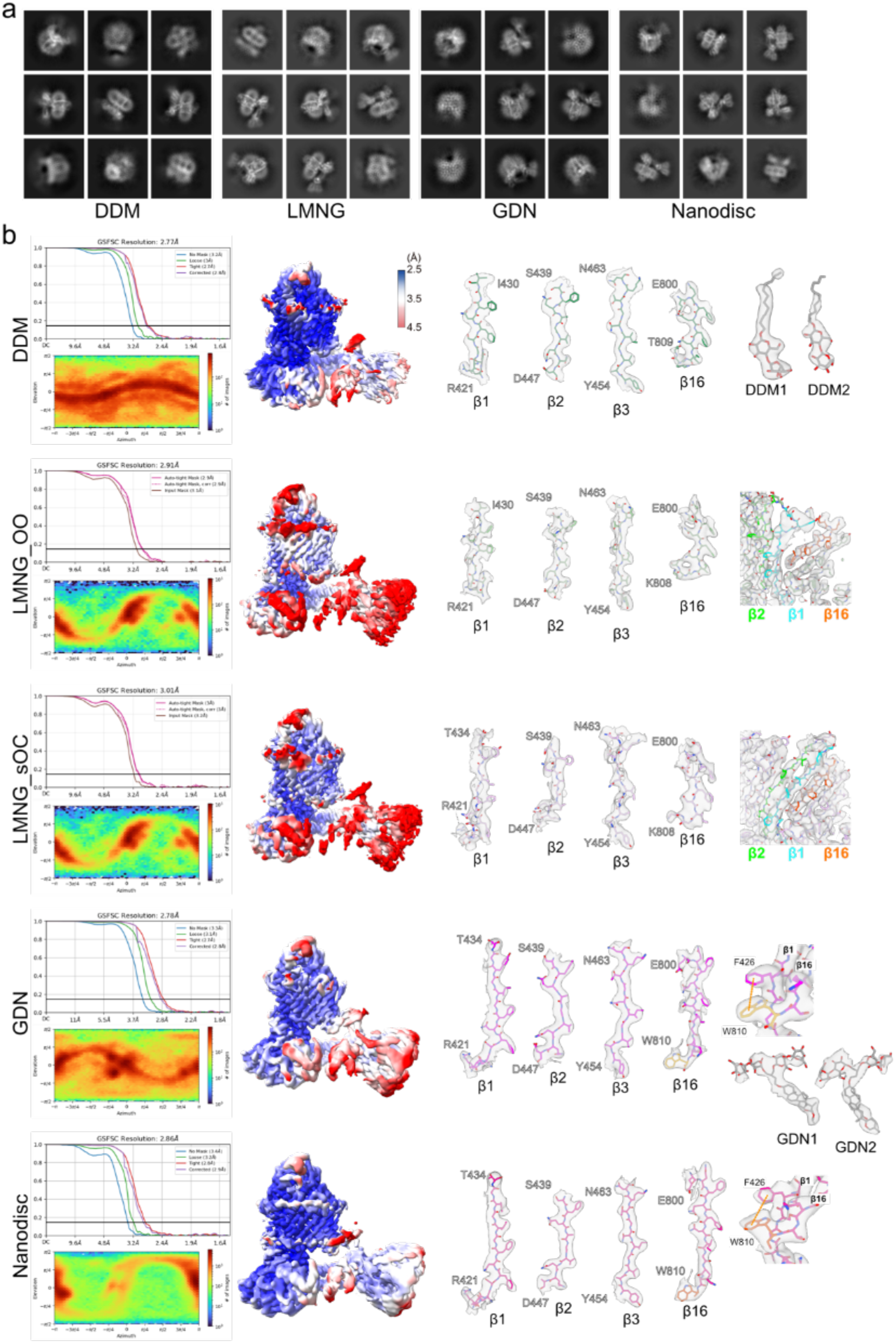
Map quality assessment. (a) Representative 2D class averages for each condition. (b) For each map, the gold-standard FSC resolution curve, the angular distribution, the local-resolution-colored map, and representative densities for the β1, β2, β3, and β16 strands are shown. Densities for bound detergent molecules are shown where present.

**Supplementary Figure 4.**
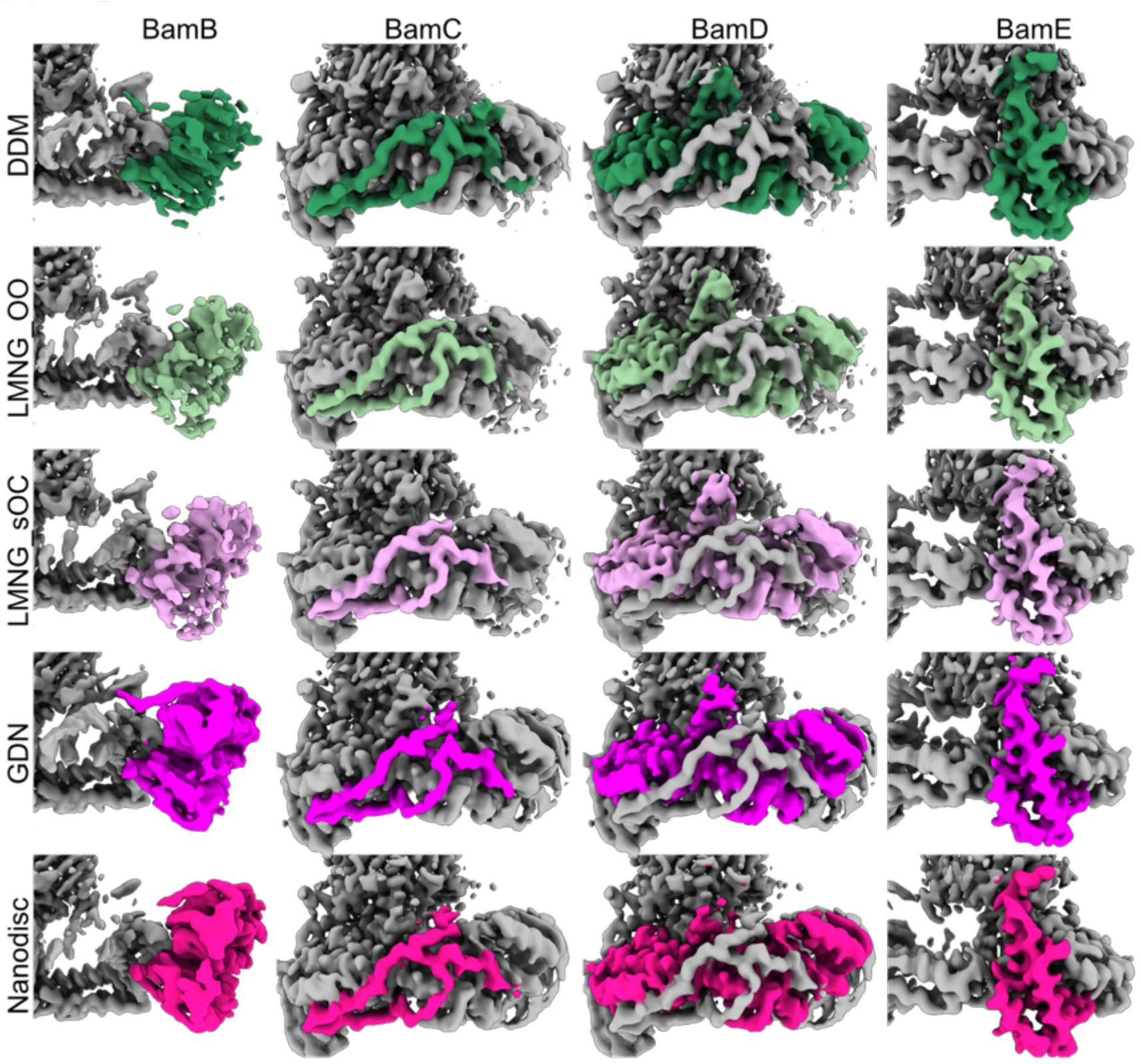
Cryo-EM densities of BAM accessory subunits under different membrane-mimetic conditions. Cryo-EM densities of BamB, BamC, BamD, and BamE (columns, left to right) in DDM, LMNG-OO, LMNG-sOC, GDN, and nanodiscs (rows, top to bottom). The density corresponding to each indicated subunit is colored, and the remainder of the complex is shown in gray. OO, outward-open state. sOC, semi-occluded state.

**Supplementary Figure 5.**
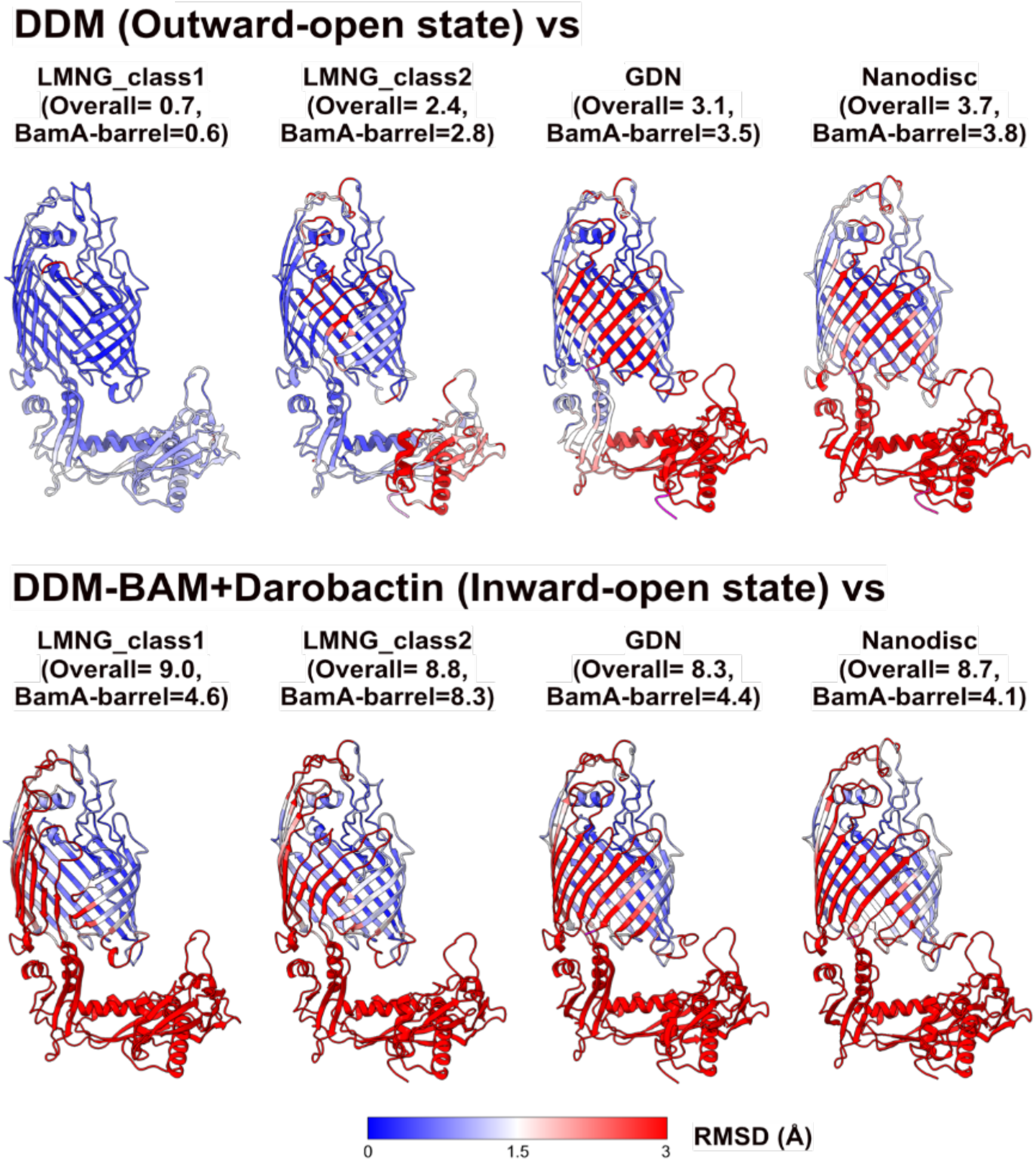
Per-residue RMSD between BamA structures. Structures are superposed and colored by per-residue Cα deviation using the color scale shown (0 to 3 Å). The top row compares DDM (outward-open) with LMNG-class1, LMNG-class2, GDN, and nanodisc. The bottom row compares the inward-open reference (DDM-BAM plus darobactin, 7NRI) with the same structures. The overall and BamA-barrel RMSD values are given above each panel.

**Supplementary Figure 6.**
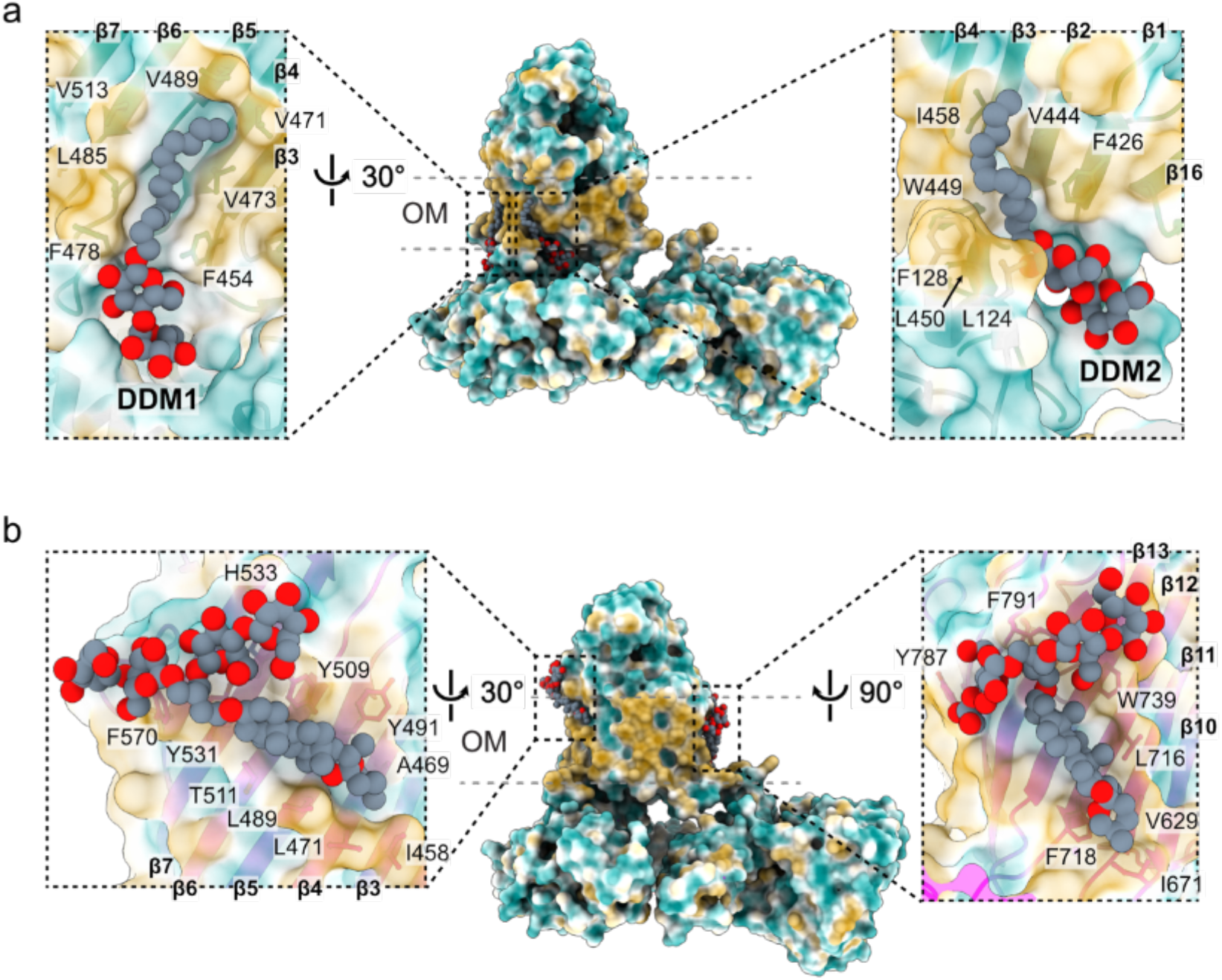
Detergent molecules modeled on the lateral surface of the BamA β-barrel. (a) Two DDM molecules, DDM1 and DDM2, were modeled in well-defined densities on the lateral surface of the BamA β-barrel within the membrane-spanning region. Enlarged views show the surrounding hydrophobic residues and β-strands contributing to the detergent-binding grooves. (b) GDN molecules were likewise modeled along the lateral surface of the BamA β-barrel in the GDN-OC structure. Enlarged views show GDN molecules occupying hydrophobic grooves formed by the transmembrane β-strands and surrounding hydrophobic residues. The approximate outer membrane boundaries are indicated by dashed lines. Views are related by the indicated rotations.

**Supplementary Figure 7.**
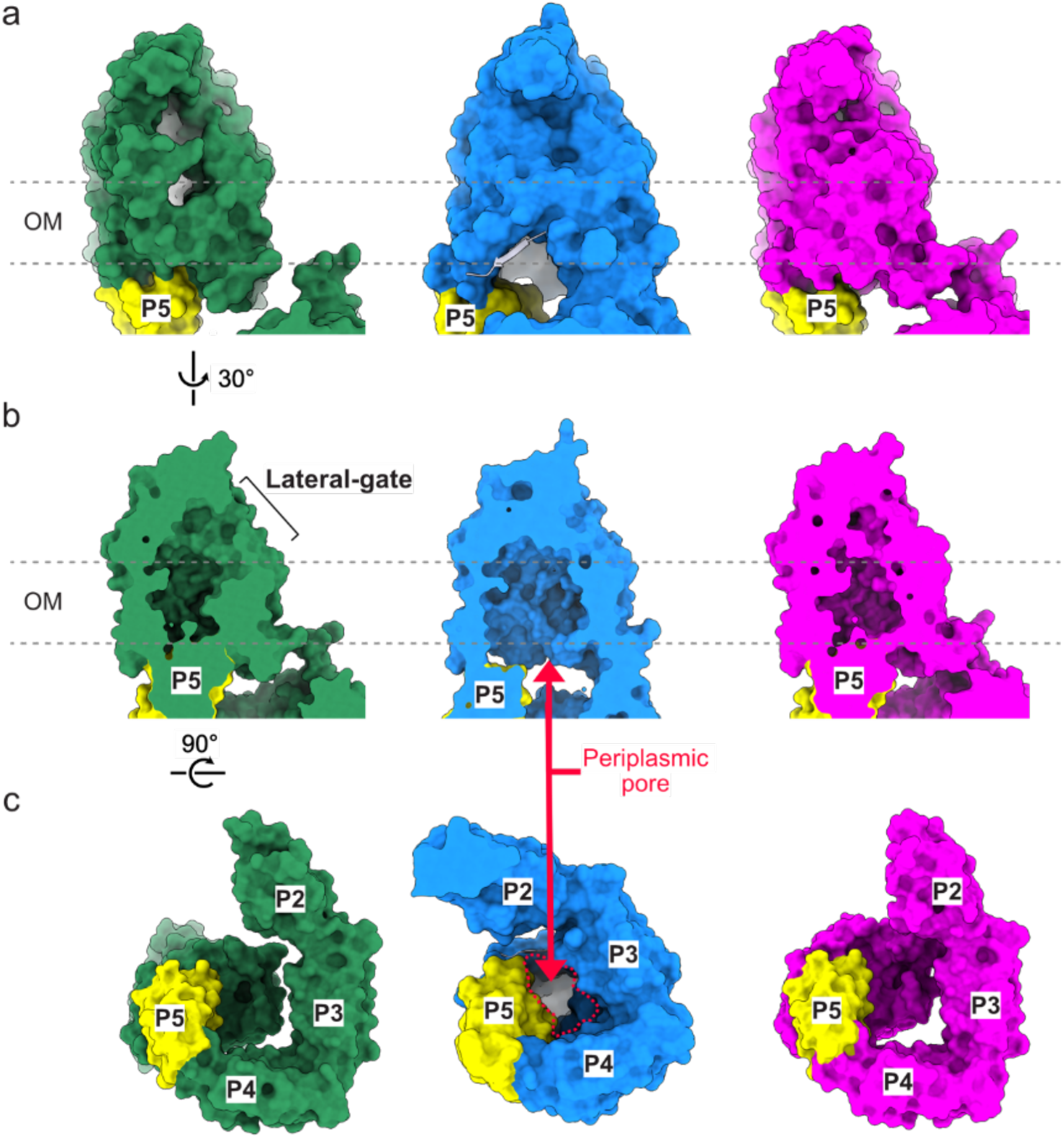
Surface representation of the lateral gate and periplasmic pore. (a) Surface views of the OO (green), IO (blue), and OC (magenta) states, with P5 highlighted and the OM indicated. (b) Cut-away views after a 30° rotation, showing the lateral gate and the periplasmic pore that is open only in the IO state (red arrow). (c) Bottom views of the POTRA ring (P2 to P5) after a further 90° rotation. The periplasmic opening seen in the IO state is closed in the OO and OC states.

**Supplementary Figure 8.**
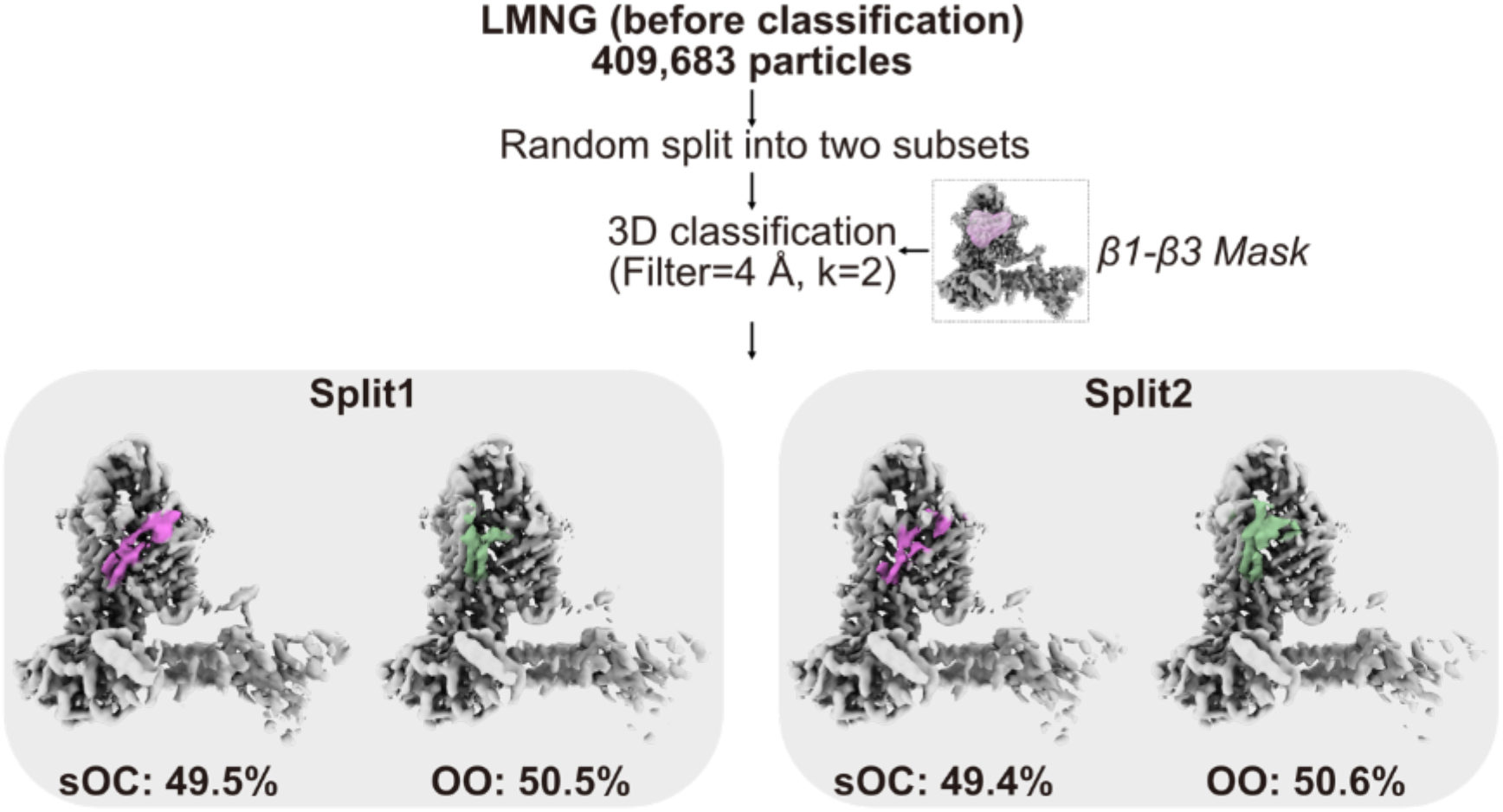
Reproducibility of the semi-occluded class. The LMNG particles were randomly split into two subsets, and focused 3D classification with a β1-β3 mask was applied to each. The OO and sOC classes were reproduced at nearly equal proportions in both subsets (sOC 49.5% and 49.4%).

**Supplementary Figure 9.**
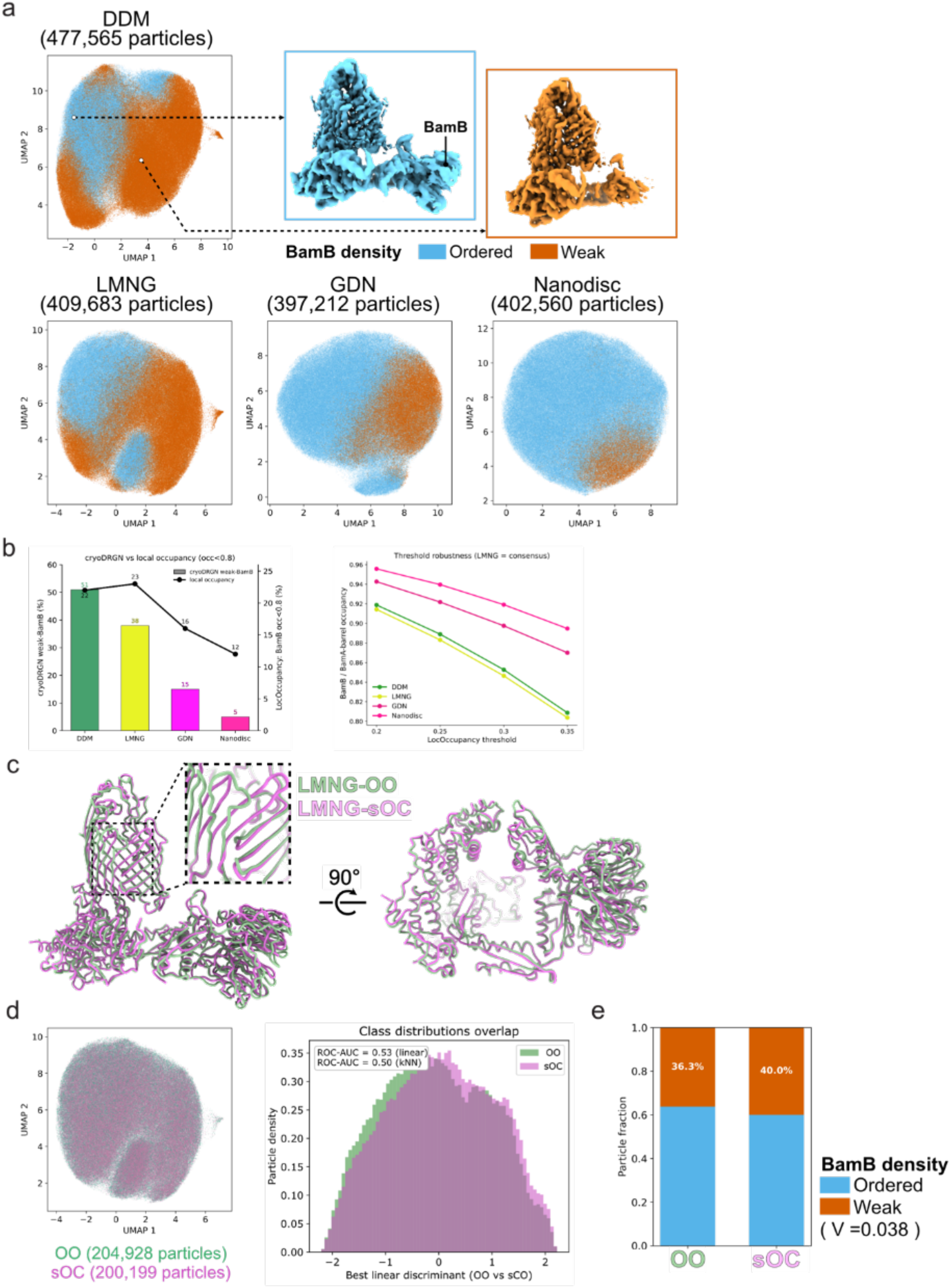
Membrane environment regulates BamB structural ordering independently of BamA lateral-gate conformation. (a) cryoDRGN UMAP latent embeddings for each condition, colored by BamB density (ordered in blue, weak in orange). Representative maps for the ordered and weak populations are shown for DDM, with the BamB position indicated. (b) Comparison of BamB structural ordering quantified by cryoDRGN and local occupancy analysis (LocOccupancy). Left, fraction of particles with weakly ordered BamB estimated by cryoDRGN (bars) and fraction of BamB residues with local occupancy below 0.8 determined by LocOccupancy (line). Right, robustness of the LocOccupancy analysis across different occupancy thresholds. The relative ordering among the four membrane-mimetic conditions was preserved over the tested threshold range. (c) Superposition of the LMNG-OO and LMNG-sOC models, with a close-up of the lateral gate, in two views related by a 90° rotation. (d) cryoDRGN latent embedding colored by the OO and sOC labels obtained from 3D classification (left) and the overlap of the two class distributions along the best linear discriminant (right). The two states are not separated (ROC-AUC 0.53 linear and 0.50 kNN). (e) Fraction of particles with weak BamB density for the OO and sOC classes, showing no meaningful difference (36.3% versus 40.0%, Cramér’s V = 0.038).

**Supplementary Figure 10.**
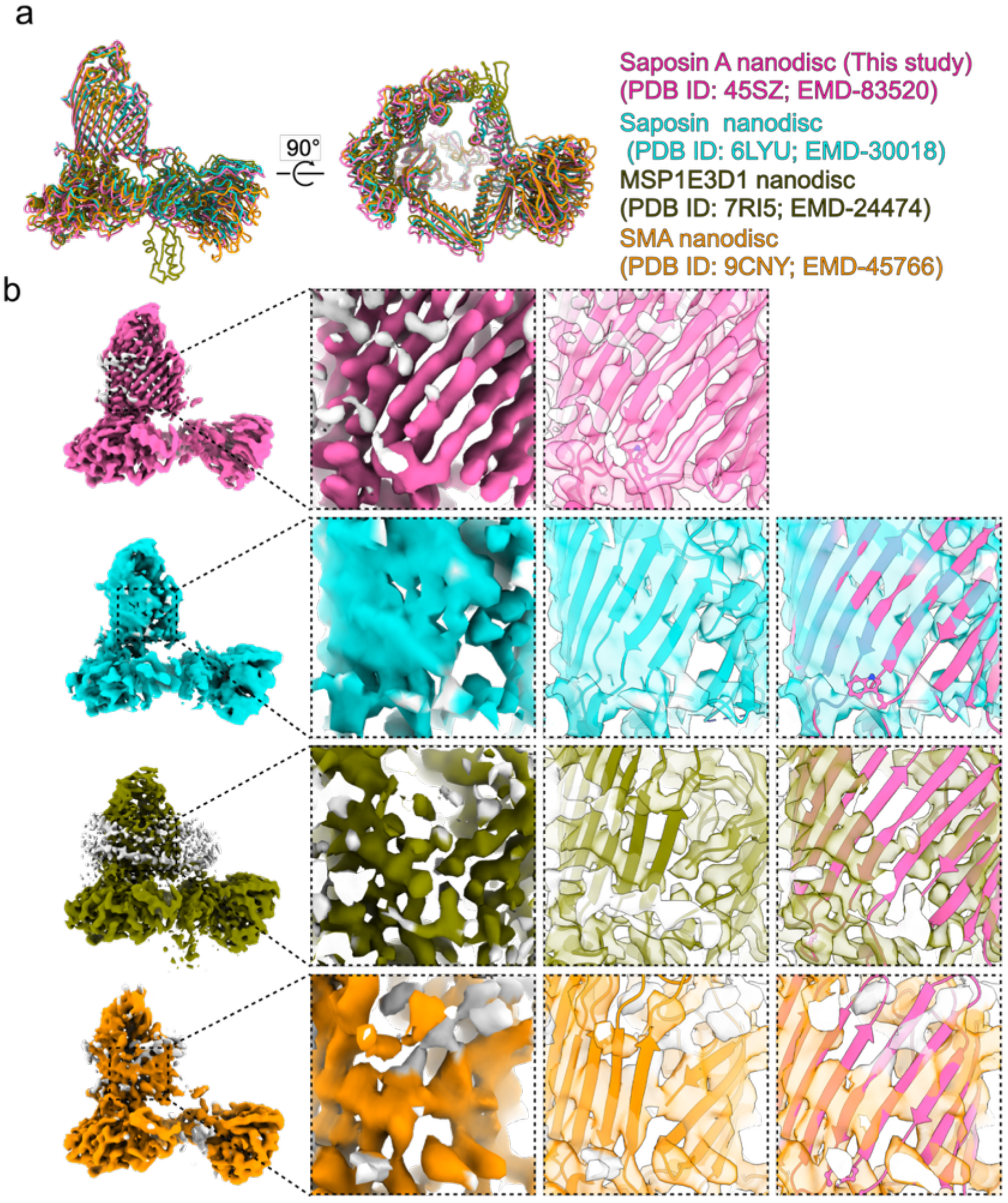
Comparison with previously reported lipid-bilayer BAM structures. (a) Superposition of Saposin A-OC (this study, PDB 45SZ, EMD-83520), Saposin nanodisc (6LYU, EMD-30018), MSP1E3D1 nanodisc (7RI5, EMD-24474), and SMA nanodisc (9CNY, EMD-45766), in two views related by a 90° rotation. (b) Gate-region density for each structure. From left to right, the map, the map with the deposited model, and an overlay with the model of this study are shown. The β-strand densities are clearly separated only in the structure of this study.

**Supplementary Figure 11.**
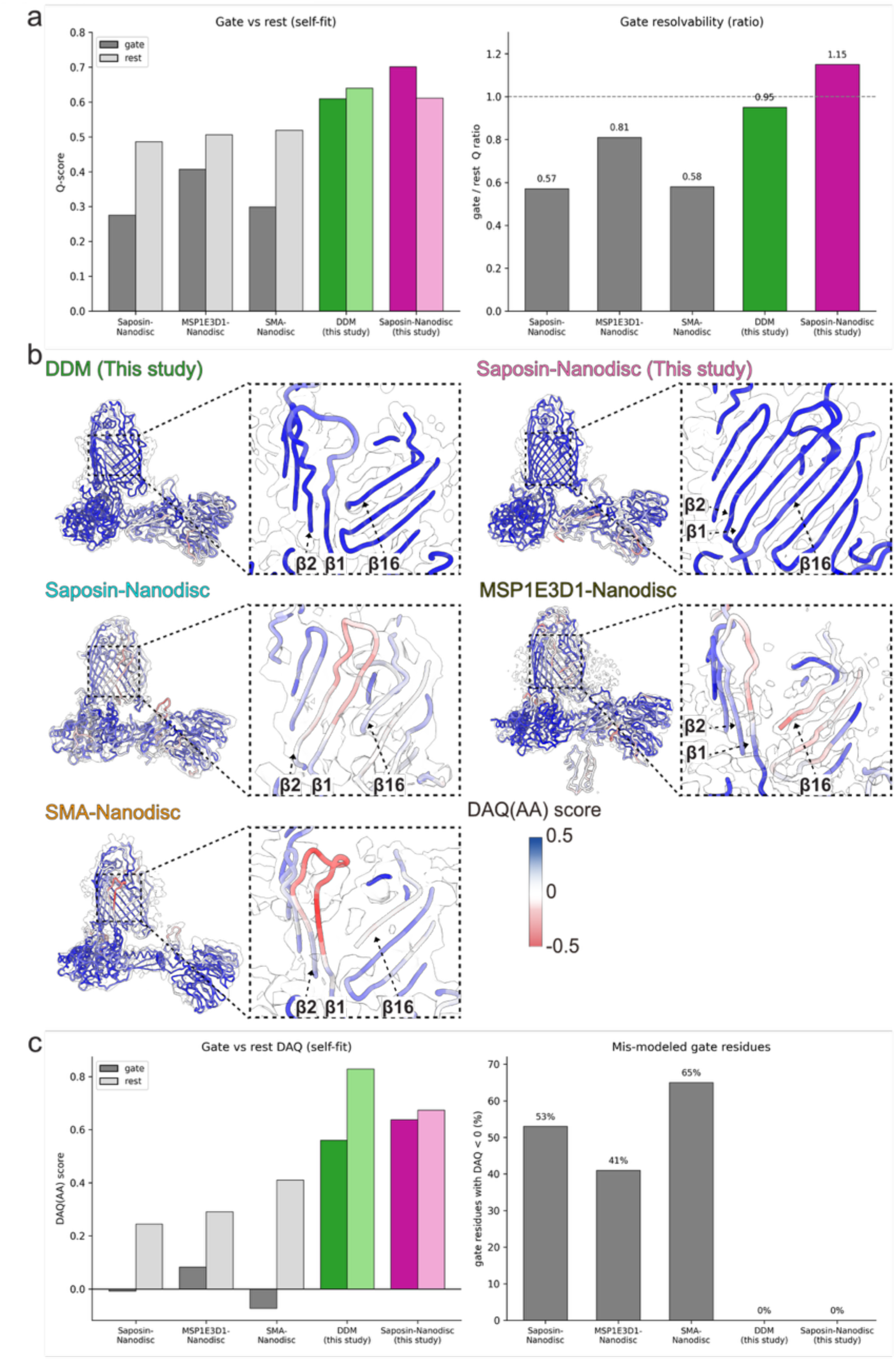
Quantitative assessment of gate map quality by Q-score and DAQ. (a) Self-fit Q-score for the gate and the rest of the model (left) and the gate-to-rest Q-score ratio (right) for Saposin-Nanodisc (6LYU), MSP1E3D1-Nanodisc (7RI5), SMA-Nanodisc (9CNY), DDM (this study), and Saposin-Nanodisc (this study). (b) Models colored by DAQ (AA) score using the scale shown, with a close-up of the gate region for each structure. (c) Self-fit DAQ (AA) for the gate and the rest of the model (left) and the fraction of gate residues with a negative DAQ score, which indicates probable mis-modeling (right).

**Supplementary Figure 12.**
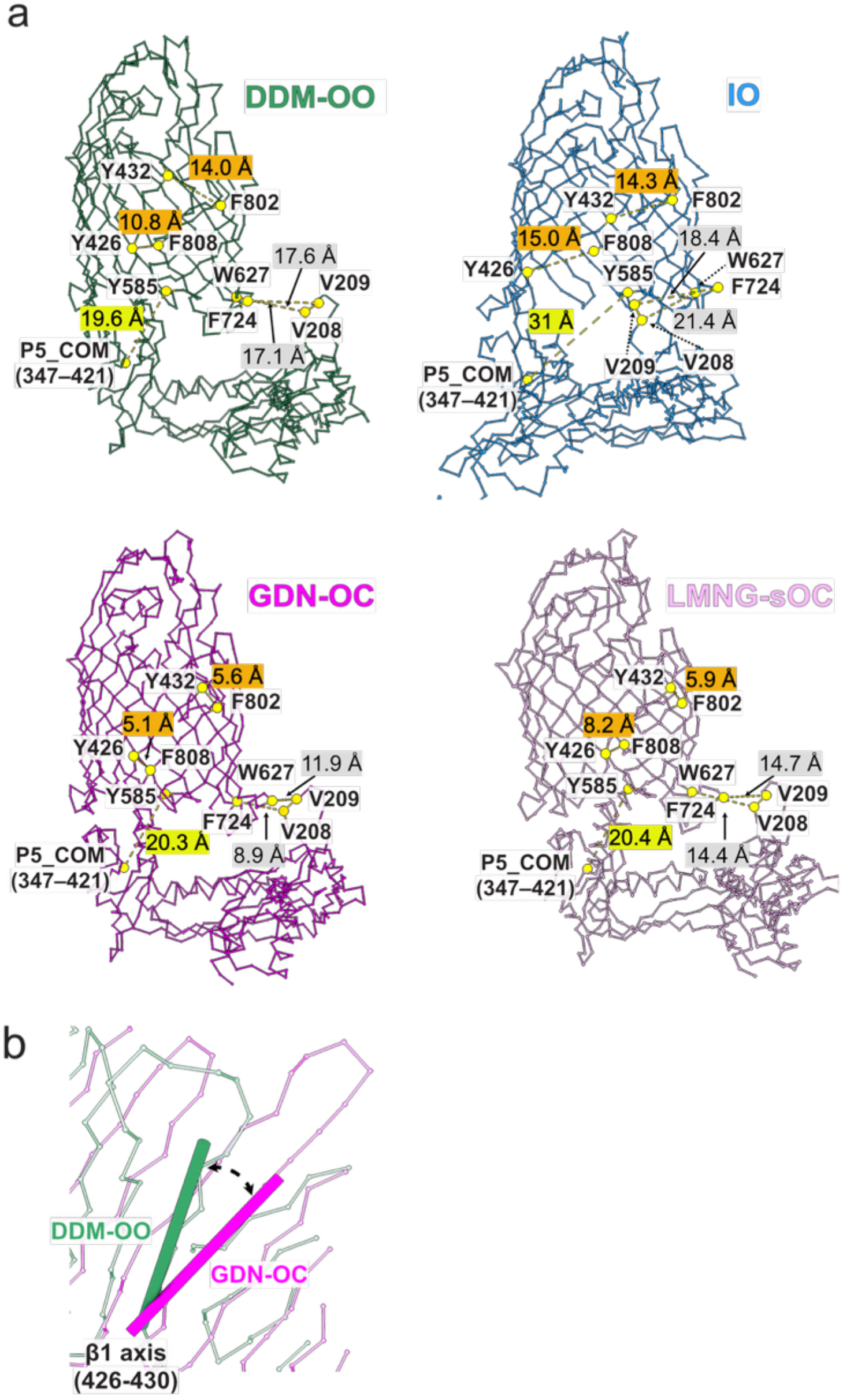
Measurement of gate geometry. (a) Distances measured in DDM-OO (green), IO (blue), GDN-OC (magenta), and LMNG-sOC (light purple). Measurements include the β1–β16 seam distances between Y432 and F802 and between Y426 and F808, the distance from Y585 to the center of mass of POTRA5 (P5, residues 347–421), and the periplasmic distances between W627 and V209 and between F724 and V208. Distances are indicated in ångströms. (b) β1 swing angle defined by the principal axes of β1 (residues 426–430) in DDM-OO and GDN-OC after superposition on the barrel core. OO, outward-open state. IO, inward-open state. OC, occluded state. sOC, semi-occluded state.Table

## Supplementary Table

**Supplementary Table 1.** Apparent relative subunit abundance of BAM under the four conditions.

| Subunit | Stain-free (Trp-corrected) |  |  |  | CBB (MW-corrected) |  |  |  |
| --- | --- | --- | --- | --- | --- | --- | --- | --- |
|  | DDM | LMNG | GDN | Nanodisc | DDM | LMNG | GDN | Nanodisc |
| BamA | 1.00 | 1.00 | 1.00 | 1.00 | 1.00 | 1.00 | 1.00 | 1.00 |
| BamB | 0.68 ± 0.05 | 0.75 ± 0.01 | 0.79 ± 0.04 | 0.83 ± 0.02 | 0.56 ± 0.04 | 0.68 ± 0.03 | 0.71 ± 0.00 | 0.60 ± 0.06 |
| BamC | 0.96 ± 0.10 | 1.01 ± 0.04 | 0.95 ± 0.05 | 0.48 ± 0.02 | 1.04 ± 0.14 | 1.07 ± 0.02 | 1.04 ± 0.02 | 0.46 ± 0.04 |
| BamD | 1.27 ± 0.10 | 1.12 ± 0.06 | 1.11 ± 0.06 | 1.21 ± 0.09 | 1.69 ± 0.21 | 1.57 ± 0.04 | 1.65 ± 0.08 | 1.55 ± 0.01 |
| BamE | 1.66 ± 0.09 | 3.02 ± 0.15 | 3.01 ± 0.13 | 0.89 ± 0.06 | 1.39 ± 0.24 | 2.79 ± 0.22 | 2.85 ± 0.40 | 0.65 ± 0.06 |

Apparent subunit abundance was quantified by stain-free detection (corrected for tryptophan content) or Coomassie Brilliant Blue staining (corrected for molecular weight), normalized to BamA (=1), and presented as the mean ± SD (N = 3). Because wild-type BamE contains no tryptophan residues, stain-free detection of BamE relied on tryptophan residues present in the affinity-tagged BamE-Strep construct.

**Supplementary Table 2.** Pairwise Cα RMSD values among BAM conformational states.

|  | DDM-OO | LMNG-OO | LMNG-sOC | GDN-OC | Nanodisc-OC | 7NRI-IO |
| --- | --- | --- | --- | --- | --- | --- |
| DDM-OO | - | 0.7 | 2.4 | 3.1 | 3.7 | 9.1 |
| LMNG-OO |  | - | 2.2 | 3.1 | 3.6 | 9.0 |
| LMNG-sOC |  |  | - | 1.8 | 2.6 | 8.8 |
| GDN-OC |  |  |  | - | 1.9 | 8.3 |
| Nanodisc-OC |  |  |  |  | - | 8.7 |
| 7NRI-IO |  |  |  |  |  | - |

matched Cα per pair ≥ 715 BamA-barrel (422-810)
|  | DDM-OO | LMNG-OO | LMNG-sOC | GDN-OC | Nanodisc-OC | 7NRI-IO |
| --- | --- | --- | --- | --- | --- | --- |
| DDM-OO | - | 0.6 | 2.8 | 3.5 | 3.8 | 4.7 |
| LMNG-OO |  | - | 2.7 | 3.5 | 3.7 | 4.6 |
| LMNG-sOC |  |  | - | 1.8 | 2.3 | 4.5 |
| GDN-OC |  |  |  | - | 1.4 | 4.4 |
| Nanodisc-OC |  |  |  |  | - | 4.1 |
| 7NRI-IO |  |  |  |  |  | - |

matched Cα per pair ≥ 387 BamB
|  | DDM-OO | LMNG-OO | LMNG-sOC | GDN-OC | Nanodisc-OC | 7NRI-IO |
| --- | --- | --- | --- | --- | --- | --- |
| DDM-OO | - | 0.5 | 1.7 | 1.6 | 1.4 | 1.5 |
| LMNG-OO |  | - | 1.7 | 1.6 | 1.3 | 1.5 |
| LMNG-sOC |  |  | - | 0.7 | 1.5 | 1.9 |
| GDN-OC |  |  |  | - | 1.5 | 1.8 |
| Nanodisc-OC |  |  |  |  | - | 1.2 |
| 7NRI-IO |  |  |  |  |  | - |

matched Cα per pair ≥ 347 BamC
|  | DDM-OO | LMNG-OO | LMNG-sOC | GDN-OC | Nanodisc-OC | 7NRI-IO |
| --- | --- | --- | --- | --- | --- | --- |
| DDM-OO | - | 0.7 | 2.6 | 3.6 | 4.2 | 4.5 |
| LMNG-OO |  | - | 2.6 | 3.2 | 4.3 | 4.2 |
| LMNG-sOC |  |  | - | 0.8 | 2.2 | 2.3 |
| GDN-OC |  |  |  | - | 2.2 | 2.3 |
| Nanodisc-OC |  |  |  |  | - | 2.7 |
| 7NRI-IO |  |  |  |  |  | - |

BamD
|  | DDM-OO | LMNG-OO | LMNG-sOC | GDN-OC | Nanodisc-OC | IO (7NRI) |
| --- | --- | --- | --- | --- | --- | --- |
| DDM-OO | - | 0.6 | 1.0 | 1.0 | 1.2 | 1.2 |
| LMNG-OO |  | - | 0.7 | 1.0 | 1.3 | 1.3 |
| LMNG-sOC |  |  | - | 0.9 | 1.3 | 1.2 |
| GDN-OC |  |  |  | - | 0.7 | 0.7 |
| Nanodisc-OC |  |  |  |  | - | 0.7 |
| 7NRI-IO |  |  |  |  |  | - |

BamE
|  | DDM-OO | LMNG-OO | LMNG-sOC | GDN-OC | Nanodisc-OC | 7NRI-IO |
| --- | --- | --- | --- | --- | --- | --- |
| DDM-OO | - | 0.4 | 1.0 | 0.9 | 0.8 | 0.6 |
| LMNG-OO |  | - | 1.0 | 1.0 | 0.9 | 0.6 |
| LMNG-sOC |  |  | - | 0.4 | 1.0 | 0.6 |
| GDN-OC |  |  |  | - | 1.0 | 0.6 |
| Nanodisc-OC |  |  |  |  | - | 0.7 |
| 7NRI-IO |  |  |  |  |  | - |

**Supplementary Table 3.**
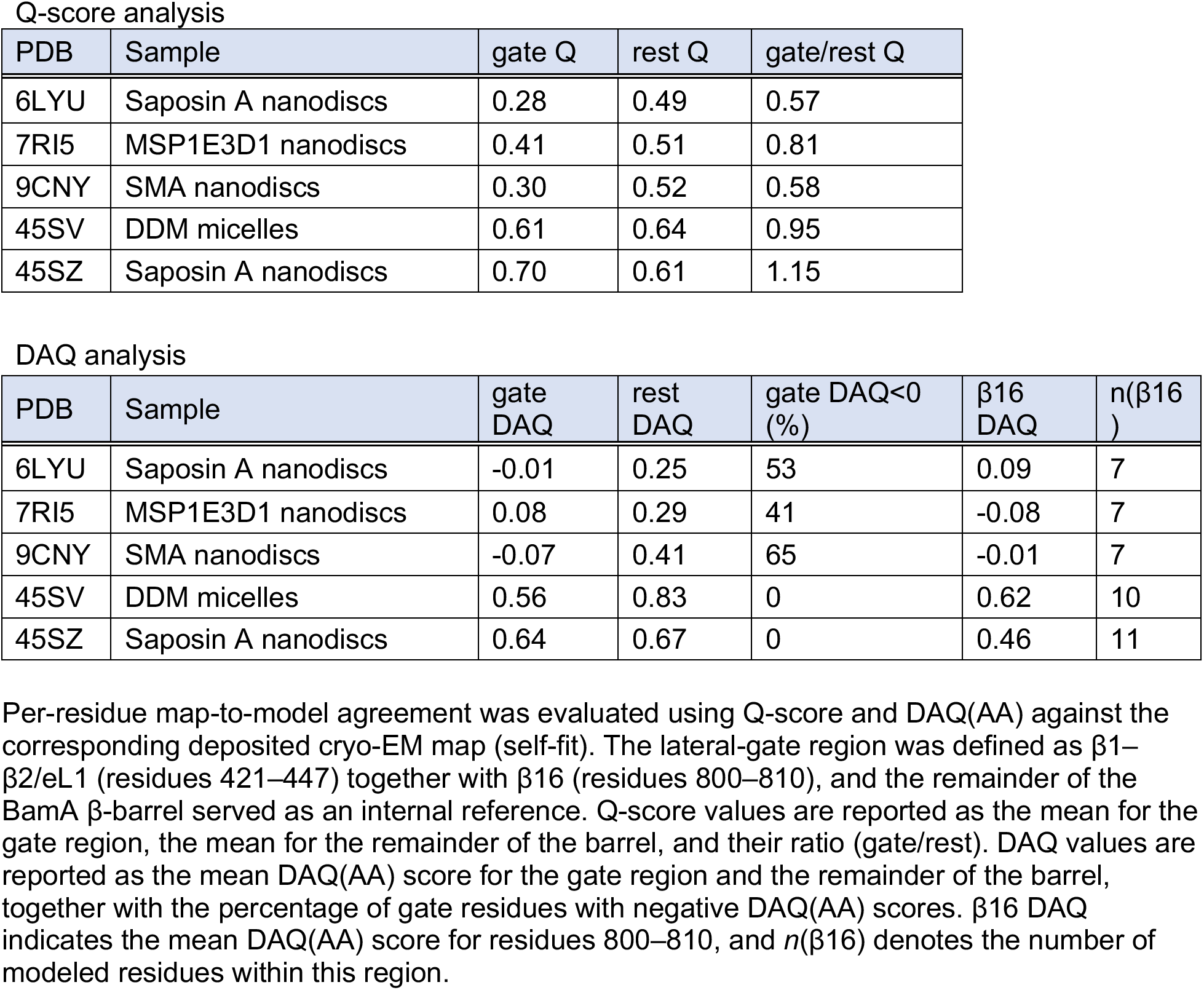
Comparison of lateral-gate map quality in previously reported and newly determined lipid-environment BAM structures.

**Supplementary Table 4.** Structural parameters describing the major BamA conformational states.

| Parameter | DDM-OO<br>(45SV) | LMNG-OO<br>(45SW) | DDM-OO<br>(9CNW) | LMNG-sOC<br>(45SX) | GDN-OC<br>(45SY) | Nanodisc-OC<br>(45SZ) | DDM-IO<br>(7NRI) |
| --- | --- | --- | --- | --- | --- | --- | --- |
| Y432–F802<br>distance (Å) | 14 | 13 | 15.3 | 5.9 | 5.6 | 5.9 | 14.3 |
| F426–K808<br>distance (Å) | 10.8 | 10.7 | 10.7 | 8.2 | 5.1 | 5.2 | 15 |
| Periplasmic<br>opening (Å) | 19.4 | 19.6 | 21.1 | 20.4 | 20.3 | 21.2 | 31 |
| T5–P3L<br>distance (Å) | 17.9 | 17.1 | 15.8 | 14.7 | 11.9 | 10 | 21.4 |
| T6–P3L<br>distance (Å) | 17.1 | 16.9 | 13.1 | 14.4 | 8.5 | 7 | 18.4 |
| β1 swing ($^{\circ}$ vs<br>OO) | 0 | 1.5 | 8.2 | 42 | 41.7 | 41.9 | 51.6 |

Distances are reported in Å. The β1– β16 seam was quantified using the Cα –Cα distances between Y432 and F802 or between F426 and K808. The periplasmic opening was measured as the distance between the Cα atom of Y585 and the center of mass (COM) of POTRA5 (residues 347–421). The T5–P3L and T6–P3L distances correspond to the Cα –Cα distances between V209 and W627 and between V208 and F724, respectively. The β1 swing angle was calculated from the principal axis of β1 (residues 426–430) relative to DDM-OO after superposition on the BamA barrel core (residues 440–795). Values are based on chain A using *E. coli* BamA residue numbering.

## Methods

### Expression and purification of the BAM

BAM samples were prepared following established protocols^4,32^ with slight modifications. *E. coli* BL21(DE3) cells (NEB) were transformed with the appropriate plasmids and grown overnight in LB medium. These cultures were used to inoculate 1 L of LB in 2 L baffled flasks (total volume: 8 L), supplemented with 100 µg/mL ampicillin. After growth at 37 °C to an OD₆₀₀ of 0.6, protein expression was induced by adding 0.1 mM IPTG, followed by overnight incubation at 20 °C. Cells were collected by centrifugation, resuspended in buffer A (50 mM Tris-HCl, pH 8.0, 150 mM NaCl, 10% glycerol), and lysed via sonication. Centrifugation at 27,000 × g for 10 min at 4 °C removed debris. The supernatant was ultracentrifuged at 145,000 × g for 1 h (Ti45 rotor, 4 °C) to isolate membrane fractions. These membranes were solubilized in buffer A containing 2% DDM and gently agitated at 4 °C. After clarification by centrifugation, the solubilized proteins were incubated with Strep-Tactin Sepharose resin (IBA) for 10 min. After loading, the resin was washed with buffer B containing 50 mM Tris-HCl, pH 8.0, and 150 mM NaCl. For the DDM sample, 20 bed volumes of buffer B containing 0.03% DDM were used. For detergent exchange, the LMNG and GDN samples were washed with 50 bed volumes of buffer B containing 0.003% LMNG or 0.003% GDN, respectively. Proteins were eluted using the same buffer supplemented with 5 mM D-desthiobiotin. The eluted BAM was concentrated using a 100 kDa MWCO centrifugal filter (Amicon Ultra) and further purified by size-exclusion chromatography on a Superose 6 Increase 10/300 GL column (GE Healthcare), except that the LMNG sample was purified using a Superose 6 Increase 3.2/300 column. Size-exclusion chromatography was performed in buffer C containing 20 mM HEPES-KOH, pH 8.0, and 150 mM NaCl, supplemented with 0.03% DDM, 0.003% LMNG, or 0.003% GDN depending on the sample. Peak fractions were pooled, concentrated to 20 mg/mL, flash-frozen in liquid nitrogen, and stored at –80 °C.

### Reconstitution in Saposin A nanodiscs

BAM was reconstituted into Saposin A nanodiscs essentially as described previously.^8^ The *E. coli* polar lipid extract (Avanti Polar Lipids) was suspended in lipid buffer (20 mM Hepes-KOH (pH 8.0), 150 mM NaCl, and 1% (w/v) DDM) at 37 °C for 1 h. Saposin A membrane scaffold protein and purified DDM-solubilized BAM were added to the reconstitution mixture at a final molar ratio of 1:12:40 (BAM: Saposin A: lipid) and incubated at 4 °C for 1 h. Detergent was removed by incubation with 100 mg/mL Bio-Beads SM-2 (Bio-Rad). The Bio-Beads were replaced with fresh Bio-Beads every 4 h for a total of three rounds of detergent removal. Following detergent removal, the nanodisc-reconstituted BAM was applied to a Superose 6 Increase 10/300 GL SEC column in buffer C and the peak fractions were collected and concentrated to 2 mg/mL.

### SDS-Page and subunit quantification

Purified BAM from each condition was analyzed by SDS-PAGE (15% polyacrylamide). The same gel was first imaged by stain-free detection^20^, which depends on a UV-induced reaction between tryptophan residues and the 2,2,2-trichloroethanol incorporated in the gel, and was subsequently stained with Coomassie Brilliant Blue (CBB). Band intensities were quantified in ImageJ^33^. For each subunit, the intensity was corrected by dividing by the number of tryptophan residues for stain-free detection or by the molecular weight for CBB, and expressed as an apparent abundance relative to BamA (set to 1). Tryptophan counts and molecular weights used for correction were BamA (13; 90.5 kDa), BamB (9; 41.9 kDa), BamC (5; 36.8 kDa), BamD (2; 27.8 kDa), and BamE-Strep (3; 16.2 kDa). Values are presented as mean ± SD from three independent preparations (*n* = 3).

### Cryo-EM sample preparation

Freshly glow-discharged Quantifoil holey carbon grids (R0.6/1, Cu, 300 mesh) were prepared using a JEC-3000FC glow-discharge system (JEOL). Three microliters of BAM in 0.03% DDM (15 mg/mL, 75.8 µM), 0.003% LMNG (6 mg/mL, 30 µM), 0.003% GDN (12 mg/mL, 60 µM), or Saposin A nanodiscs (0.5 mg/mL, 2.5 µM) was applied to the grids. Grids were blotted for 2–3 s at a blot force of 12 and plunge-frozen in liquid ethane using a Vitrobot Mark IV (Thermo Fisher Scientific) operated at 8 °C and 99% relative humidity.

### Cryo-EM data collection

Grid quality was assessed using a Glacios (Thermo Fisher Scientific) operated at an acceleration voltage of 200 kV. Fina data for the DDM-, LMNG-, and GDN-solubilized BAM samples were collected on a CRYO ARM 300 microscope (JEOL) operated at 300 kV and equipped with a cold field-emission gun, an in-column Ω energy filter, and a K3 direct electron detector (Gatan) operated in CDS mode. Images were recorded at a nominal magnification of ×60,000, corresponding to a calibrated pixel size of 0.752 Å. Movies were recorded for 2.5 s and fractionated into 50 frames, yielding a total exposure of 50 e⁻/Å². Automated data collection was performed using SerialEM^34,35^ with image shift at a defocus range of −1.2 to −1.6 µm. Approximately 7,000–9,000 movies were collected for each dataset.

Final data for the Saposin A nanodisc sample were collected on a Titan Krios G1 (Thermo Fisher Scientific) operated at 300 kV and equipped with a K3 direct electron detector (Gatan). Images were recorded at a nominal magnification of ×60,000, corresponding to a calibrated pixel size of 0.864 Å. Movies were recorded for 2.5 s and fractionated into 50 frames, yielding a total exposure of 50 e⁻/Å². Automated data collection was performed using SerialEM^34,35^ with image shift at a defocus range of −1.2 to −1.6 µm. The number of movies collected for each dataset is summarized in Table 1.

### Cryo-EM data processing

Cryo-EM data were processed in CryoSPARC^36^ (Figure S2). Initial processing was performed using CryoSPARC v4.0.0, and subsequent refinement and reprocessing steps were carried out using later versions up to v5.0.6. Movie frames were corrected for beam-induced motion using Patch Motion Correction, followed by CTF estimation using Patch CTF Estimation. Micrographs with estimated CTF fit resolutions poorer than 5 Å were excluded using Curate Exposures.

Following curation, 8,361 DDM, 8,897 LMNG, 7,105 GDN, and 7,742 nanodisc micrographs were retained for further processing. Particles were identified by template-based picking using a particle diameter of 180 Å and were inspected using Inspect Picks. Particles were initially extracted into 64-pixel boxes for particle cleaning. This yielded 4,798,705 DDM, 5,010,408 LMNG, 3,799,021 GDN, and 5,894,167 nanodisc particles. Extracted particles were subjected to two successive rounds of reference-free 2D classification with 200 classes per round. Classes displaying well-defined BAM features were retained, resulting in 708,638 DDM, 1,489,043 LMNG, 815,966 GDN, and 1,940,942 nanodisc particles. Selected particles were subjected to four-class *Ab*-initio reconstruction followed by iterative heterogeneous refinement using four references to remove contaminants, incomplete complexes, and poorly resolved particle populations. Particles corresponding to intact BAM were retained, yielding 477,565 DDM, 409,683 LMNG, 397,212 GDN, and 805,127 nanodisc particles. After heterogeneous refinement, particles were re-extracted into 256-pixel boxes for high-resolution refinement. The selected particles were subjected to reference-based motion correction and non-uniform refinement. Density corresponding to the detergent micelle or nanodisc was then removed by particle subtraction using a mask encompassing the BAM while excluding the surrounding membrane-mimetic density. The subtracted particle stacks were subjected to local CTF refinement followed by local refinement using default parameters. Final map resolutions were estimated using the gold-standard Fourier shell correlation criterion at FSC = 0.143.

### Focused 3D classification of the lateral gate (LMNG)

To resolve lateral-gate heterogeneity in the LMNG dataset, the consensus particle set of 409,683 particles was subjected to focused 3D classification without alignment in CryoSPARC using a mask encompassing the β1–β3 region of the BamA barrel with a filter resolution of 4 Å and *k* = 2. This analysis yielded an outward-open class and a semi-occluded class. During classification and subsequent local refinement, a total of 4,556 particles were excluded. The final LMNG-OO and LMNG-sOC particle sets contained 204,928 and 200,199 particles, respectively, and yielded maps at final resolutions of 2.91 Å and 3.01 Å..

To assess the robustness of the classification, the particle set was randomly divided into two halves and each subset was classified independently under identical conditions. Both subsets reproducibly yielded OO and sOC classes with nearly identical class distributions (sOC, 49.5% and 49.4%).

### Model building and refinement

Initial atomic models were generated by rigid-body fitting of previously reported *E. coli* BAM structures into the cryo-EM maps. The outward-open BAM structure (PDB 7ND0^22^) was used as the initial model for the DDM-OO and LMNG-OO reconstructions, whereas the laterally closed BAM structure (PDB 6LYU^8^) was used as the initial model for the LMNG-sOC, GDN-OC, and Nanodisc-OC reconstructions. Models were manually rebuilt in Coot^37,38^. DDM and GDN molecules were modeled where supported by the density maps, and the C-terminal region of β16, including W810, was built when clear density was present. Real-space refinement was performed using Phenix^39,40^ with secondary-structure and Ramachandran restraints, and the final models were validated using MolProbity^41^. Data collection, reconstruction, and refinement statistics are summarized in Table 1.

### Heterogeneity analysis (cryoDRGN)

Conformational heterogeneity was analyzed using cryoDRGN v3.5.3^23^. Particle stacks used for the final local refinement in cryoSPARC, together with the corresponding CTF parameters and particle poses, were used as input. The DDM, LMNG, and GDN datasets contained 477,565, 409,683, and 397,212 particles, respectively. The final nanodisc reconstruction contained 805,127 particles. Because this dataset was substantially larger than the detergent datasets, 402,560 particles were randomly selected to reduce differences in dataset size among conditions. No structural criterion was used for this subsampling. Particle images were downsampled to a box size of 128 × 128 pixels, corresponding to pixel sizes of 1.50–1.73 Å depending on the dataset. A circular real-space window with a radius of 0.85 was applied. Particle poses obtained from cryoSPARC refinement were kept fixed during training and were not further optimized by pose SGD. A variational autoencoder with an eight-dimensional latent variable was trained independently for each dataset for 50 epochs. The network consisted of a three-layer residual encoder and a three-layer decoder with 256 hidden units per layer. ReLU activation, Gaussian positional encoding, and a Fourier-domain representation were used. Models were trained using the Adam optimizer with a learning rate of 1 × 10^-4^, a batch size of 16, and no weight decay. The weight applied to the Kullback–Leibler divergence term was 0.125. Mixed-precision training was enabled. The same network architecture and training parameters were used for all four datasets. The resulting eight-dimensional latent coordinates were visualized in two dimensions using UMAP^24^. UMAP was used for visualization and was not used for quantitative clustering. To assess BamB structural ordering, the eight-dimensional latent space of each dataset was partitioned into 20 clusters by k-means clustering. Clustering was performed using the cryoDRGN implementation with scikit-learn v1.7.2. K-means++ initialization was used with a single initialization, a fixed random state of 0, and a maximum of 10 iterations. For each k-means centroid, the nearest observed particle latent coordinate was selected as the representative point, and a three-dimensional volume was generated from this coordinate using the trained cryoDRGN decoder. Representative volumes were generated without additional low-pass filtering, cropping, or downsampling. The 20 representative volumes from each dataset were visually inspected at the expected position of BamB. Volumes showing continuous and clearly resolved BamB density were classified as well ordered, whereas volumes showing markedly attenuated, fragmented, or largely absent BamB density were classified as weakly ordered. Each particle was assigned according to the classification of its corresponding k-means cluster, and the fraction of particles associated with weakly ordered BamB was calculated for each membrane-mimetic condition.

To examine the relationship between BamA lateral-gate conformation and BamB ordering in the LMNG dataset, particles assigned to the OO and sOC classes by focused three-dimensional classification in CryoSPARC were mapped onto their corresponding positions in the cryoDRGN latent space. The separability of the OO and sOC populations was evaluated using receiver operating characteristic analysis based on linear discriminant analysis and a k-nearest-neighbor classifier. Association between the BamA gate state and BamB ordering was independently evaluated using Cramér’s V.

### Local occupancy analysis

Local electron density occupancy maps were computed with LocOccupancy from the LocSpiral suite^25^. For each condition the unfiltered (unsharpened) map and a binary mask of the complex were used as input, with a pixel size of 0.752 Å for the DDM, LMNG and GDN maps and 0.864 Å for the Saposin A-nanodisc map, a low-resolution limit of 15 Å, a high-resolution limit of 2.8 Å, an occupancy threshold of 0.25 and a bandwidth of 5. For LMNG the consensus map from all 409,683 particles (2.67 Å) was used so that the analysis matched the cryoDRGN analysis, which pools the same particles. Local occupancy values were read at the atomic coordinates (nearest voxel) of BamB (chain B) and of the BamA barrel (chain A, residues 440 to 810). BamB ordering was quantified as the fraction of BamB residues with a local occupancy below 0.8. The BamA barrel served as an internal control, and its median local occupancy was at least ∼0.98 in every condition, confirming that map quality was comparable across conditions. The ranking of the conditions was preserved across occupancy thresholds from 0.20 to 0.35 (Figure S9).

### In vitro OMP folding assay with OmpT

The assay was performed essentially according to previously described methods^3,18,42^, with minor modifications. OmpT and the periplasmic chaperone SurA were expressed and purified from *E. coli* as previously described^42,43^. Two reaction mixtures were prepared in 20 mM Tris-HCl (pH 6.5). Mixture A (25 µL) contained 0.2 µM BAM reconstituted into Saposin A nanodiscs, 100 µM fluorogenic substrate Cy3-ARRAK-Cy5, and 0.1 mg/mL *E. coli* total lipids. Empty nanodiscs lacking BAM were prepared in parallel as negative controls. A fluorogenic peptide, Cy3-ARRAK-Cy5, was synthesized by GenScript. Mixture B (25 µL) contained 20 µM urea-denatured OmpT and 140 µM SurA. Mixture B was pre-incubated at 25 °C for 10 min and then mixed 1:1 with Mixture A to initiate OmpT folding. After mixing, reactions were incubated for an additional 10 min at 25 °C before fluorescence measurements. Fluorescence was monitored using a Varioskan LUX microplate reader (Thermo Fisher Scientific) with excitation at 490 nm and emission at 512 nm for 60 min at 30 s intervals. Fluorescence traces were baseline-corrected by subtracting the signal at *t* = 0 s. Unless otherwise stated, data represent three independent experiments (*n* = 3).

### Comparison with previously reported BAM structures determined in lipid environments

Deposited cryo-EM maps and atomic models of previously reported *E. coli* BAM structures determined in lipid bilayers (PDB 6LYU/EMD-30018, PDB 7RI5/EMD-24474, and PDB 9CNY/EMD-45766) were retrieved from the Protein Data Bank (PDB) and Electron Microscopy Data Bank (EMDB) and compared with the DDM-OO and Nanodisc-OC structures determined in this study. For each structure, per-residue map-to-model agreement was evaluated using the Q-score calculated with the QScore^26^ plugin in UCSF ChimeraX^44,45^ and the DAQ(AA) score calculated with the DAQ program^27,46^ against the corresponding deposited cryo-EM map (self-fit). The lateral-gate region was defined as β1–β2/eL1 (residues 421–447) and β16 (residues 800–810), whereas the remaining barrel residues served as an internal reference. Local map quality around the lateral gate was quantified as the ratio of the mean Q-score for the gate region to that for the remainder of the barrel (gate-to-rest ratio) and as the fraction of gate residues with negative DAQ(AA) scores.

### Structural and geometric analysis

Structural comparisons were performed using in-house Python scripts based on **gemmi**^47^. Structures were superposed by iterative least-squares fitting using the BamA barrel core (Cα atoms of residues 440–795). Lateral-gate geometry was quantified by the β1–β16 seam distances (Cα–Cα distances between Y432–F802 and F426–K808), the β1 swing angle (defined as the angle between the principal axes of the β1 Cα atoms, residues 426–430, relative to DDM-OO), and the periplasmic opening (distance between the Cα atom of Y585 and the center of mass of POTRA5, residues 347–421). Backbone hydrogen bonds across the β1–β16 seam were identified as main-chain N–O pairs between β1 (residues 421–437) and β16 (residues 795–810) with an N–O distance of less than 3.5 Å. Model completeness was evaluated from the number of modeled residues in chain A and within the β16 C-terminal region (residues 795–810), including whether W810 was modeled. Molecular graphics were prepared using UCSF ChimeraX^44,45^ .

## RESOURCE AVAILABILITY

### Lead contact

Requests for further information and resources should be directed to and will be fulfilled by the lead contact, Tsuyoshi Imasaki.

## Materials availability

All unique reagents generated in this study are available from the lead contact upon reasonable request.

## Data and code availability

- Cryo-EM density maps and atomic coordinates have been deposited in the Electron Microscopy Data Bank (EMDB) and the Protein Data Bank (PDB) under accession codes EMD-83516 and PDB: 45SV (DDM-OO), EMD-83517 and PDB: 45SW (LMNG-OO), EMD-83518 and PDB: 45SX (LMNG-sOC), EMD-83519 and PDB: 45SY (GDN-OC), and EMD-83520 and PDB: 45SZ (Nanodisc-OC), and are publicly available as of the date of publication. Accession numbers are also listed in the key resources table.
- This paper does not report original code.
- Any additional information required to reanalyze the data reported in this paper is available from the lead contact upon request.

## Acknowledgements

We are grateful to Dr. Harris D. Bernstein for providing the plasmids used for expression of the *E. coli* BAM. We thank the staff of the cryo-EM facility at the RIKEN SPring-8 Center for support with grid screening and data collection on the EM01CT (CRYO ARM 300), EM03CT (Titan Krios G1), and EM04CT (Glacios) instruments. We thank K. Chin for research management support, and the members of the Nitta laboratory for helpful discussions. This work was supported by Japan Society for the Promotion of Science (JSPS) KAKENHI grants (JP21H05254, JP23K24057 to R.N.; 25H02379 to T. I.; JP25K22521 to K. T. 24K18062 to T. K.); JST Moonshot Research and Development Program grant (JPMJMS2024-7 to R.N.); JST FOREST (JPMJFR214K to T. I); JST CREAT (JPMJCR26T2 to K. T.); Japan Agency for Medical Research and Development (AMED) CREST grant (JP21gm161003 to T. I.); the Hyogo Science and Technology Association to R. N. The cryo-EM experiments were performed at SPring-8 with the approval of the Japan Synchrotron Radiation Research Institute (JASRI) (Proposal No. 2024B2531, 2025B2529). This research was partially supported by Research Support Project for Life Science and Drug Discovery (Basis for Supporting Innovative Drug Discovery and Life Science Research (BINDS)) from AMED under Grant Number JP22ama121001, JP23ama121001 .

## Author Contributions

Conceptualization, TK, TI; Data curation, TK; Methodology, TK, TI; Investigation, TK; Formal analysis, TK; Visualization, TK; Writing – original draft, TK; Writing – review & editing, TK, HS, MS, KT, RN and TI; Funding acquisition, MS, KT, RN, TI; Supervision, HS, TI, RN.

## Declaration of generative AI and AI-assisted technologies in the writing process

During the preparation of this work, the authors used ChatGPT and Claude to improve phrasing. After using these tools, the authors reviewed and edited the content as needed and take full responsibility for the content of the published article.

